# Global identification of RsmA/N binding sites in *Pseudomonas aeruginosa* by *in vivo* UV CLIP-seq

**DOI:** 10.1101/2020.08.24.265819

**Authors:** Kotaro Chihara, Lars Barquist, Kenichi Takasugi, Naohiro Noda, Satoshi Tsuneda

**Affiliations:** Department of Life Science and Medical Bioscience, Waseda University, Tokyo, Japan; Biomedical Research Institute, National Institute of Advanced Industrial Science and Technology (AIST), Ibaraki, Japan; Helmholtz Institute for RNA-based Infection Research (HIRI), Helmholtz Center for Infection Research (HZI), Würzburg, Germany; Faculty of Medicine, University of Würzburg, Würzburg, Germany

**Author notes:** Address correspondence to Naohiro Noda, or Satoshi Tsuneda,. Helmholtz Institute for RNA-based Infection Research (HIRI), Helmholtz Center for Infection Research (HZI), Würzburg, Germany.

## Abstract

Posttranscriptional regulation of gene expression in bacteria is performed by a complex and hierarchical signaling cascade. *Pseudomonas aeruginosa* harbors two redundant RNA-binding proteins RsmA/RsmN (RsmA/N), which play a critical role in balancing acute and chronic infections. However, *in vivo* binding sites on target transcripts and the overall impact on the physiology remains unclear. In this study, we applied *in vivo* UV crosslinking immunoprecipitation followed by RNA-sequencing (UV CLIP-seq) to detect RsmA/N binding sites at single-nucleotide resolution and mapped more than 500 peaks to approximately 400 genes directly bound by RsmA/N in *P. aeruginosa*. This also demonstrated the ANGGA sequence in apical loops skewed towards 5’UTRs as a consensus motif for RsmA/N binding. Genetic analysis combined with CLIP-seq results identified previously unrecognized RsmA/N targets involved in LPS modification. Moreover, the small non-coding RNAs RsmY/RsmZ, which sequester RsmA/N away from target mRNAs, are positively regulated by the RsmA/N-mediated translational repression of *hptB*, encoding a histidine phosphotransfer protein, and *cafA*, encoding a cytoplasmic axial filament protein, thus providing a possible mechanistic explanation for homeostasis of the Rsm system. Our findings present the global RsmA/N-RNA interaction network that exerts pleiotropic effects on gene expression in *P. aeruginosa*.

**IMPORTANCE:** The ubiquitous bacterium *Pseudomonas aeruginosa* is notorious as an opportunistic pathogen causing life-threatening acute and chronic infections in immunocompromised patients. *P. aeruginosa* infection processes are governed by two major gene regulatory systems, namely, the GacA/GacS (GacAS) two-component system and the RNA-binding proteins RsmA/RsmN (RsmA/N). RsmA/N basically function as a translational repressor or activator directly by competing with the ribosome. In this study, we identified hundreds of RsmA/N regulatory target RNAs and the consensus motifs for RsmA/N bindings by UV crosslinking *in vivo*. Moreover, our CLIP-seq revealed that RsmA/N posttranscriptionally regulate cell wall organization and exert feedback control on GacAS-RsmA/N systems. Many genes including small regulatory RNAs identified in this study are attractive targets for further elucidating the regulatory mechanisms of RsmA/N in *P. aeruginosa*.

## INTRODUCTION

The gram-negative bacterium *Pseudomonas aeruginosa* is an opportunistic pathogen infecting humans that thrives in diverse environments. It can cause serious biofilm-associated infections in burn wounds, indwelling devices, and the lungs of immunocompromised cystic fibrosis patients (1). The infection processes are governed via complex posttranscriptional regulatory mechanisms stimulated by certain environmental stresses such as nutrient starvation and antibiotic exposure (2). Such posttranscriptional control networks are typically composed of globally acting RNA-binding proteins (RBPs) together with small non-coding RNAs (sRNAs). Considering the presence of a large number of putative sRNAs and widely conserved RBPs in *P. aeruginosa*, a number of them might be incorporated into the regulatory networks for the processes.

<u>H</u>ost <u>f</u>actor protein for the replication of phage <u>q</u>β (Hfq) is one of the most well-known RBPs that acts as an RNA chaperone and helps sRNAs basepair with target mRNAs for repression or activation of the translations (3). Similar to *Escherichia coli* or *Salmonella*, Hfq contributes to varied phenotypes such as quorum sensing, virulence, and antibiotic resistance in *P. aeruginosa* (4–6). Likewise, the RNA chaperone CsrA family, originally discovered as a carbon storage regulator in *E. coli*, also plays a global role in posttranscription in many bacteria (7). Unlike Hfq, the CsrA family functions by repressing translation directly by competing with the ribosome rather than acting as a matchmaker between sRNA and mRNA in Gram-negative bacteria. CsrA family proteins bind to target RNAs via a stem-loop structure comprising an ANGGA motif. Conversely, translational suppression is inhibited by at least three sRNAs, namely, CrsB, CrsC, and McaS, each containing multiple GGA motifs in *E. coli* (8, 9).

*P. aeruginosa* has two CsrA homologs RsmA and RsmN (RsmA/N) and four redundant sRNAs RsmV, RsmW, RsmY, and RsmZ that function as antagonists of RsmA/N (10–12). RsmA and RsmN are structurally distinct with respect to the position of an α helix constituent; RsmA is composed of five consecutive β sheets and a C-terminal α helix whereas RsmN consists of an uniquely inserted α helix between the β2 and β3 sheets (13). RsmA/N-titrating sRNAs, RsmY/RsmZ (RsmY/Z), are regulated by GacA/GacS (GacAS) two-component system. When the transmembrane histidine protein kinase (HPK) GacS is activated by environmental signals involved in the transition to the stationary phase, it phosphorylates its cognate response regulator GacA (14). Upon phosphorylation, GacA promotes the transcription of RsmY/Z (15). While RsmW is activated in the stationary phase or by high temperature (11), the level of RsmV expression is relatively low during the whole course of the growth, independent of GacAS activity (12). Both RsmA/N bind to some, but not all, common regulatory targets via a conserved arginine residue. Most of the known RsmA/N targets such as genes encoding for type VI secretion systems (T6SS) and exopolysaccharide biosynthesis are subject to direct translational repression (16, 17). In contrast, the genes encoding motility and type III secretion systems (T3SS) are positively regulated by RsmA/N (14, 18), thus modulating the transition between acute and chronic infections in *P. aeruginosa*.

These relationships between RsmA/N and target genes have been extensively investigated by phenotypic assays combined with genetic analysis *in vitro*. However, the complexity of *in vivo* RNA-based regulations makes attaining a comprehensive understanding of the mode of actions of RsmA/N difficult. To address this, recent studies have identified target transcripts, that are bound by members of the CsrA/Rsm family via co-immunoprecipitation approaches (19, 20). Such methods can identify major RNA ligands. However, they do not provide sufficient information with respect to the RBPs binding sites within each transcript. Herein, we performed UV crosslinking immunoprecipitation followed by RNA-seq (UV CLIP-seq) with RsmA/N to identify their regulatory targets at a single-nucleotide resolution *in vivo*. Our approach found more than 500 potential binding sites genome-wide, which enabled us to gain new insights into the biological functions and the similarities and differences between the two redundant Rsm RBPs in *P. aeruginosa*.

## RESULTS

### Transcriptome-wide mapping of RsmA/N binding sites

As a preliminary investigation, western blotting analysis was performed using *P. aeruginosa* PAO1 chromosomal FLAG tagged strains: *rsmA::3×FLAG, rsmN*::3×FLAG, and *rsmN::3×FLAG*Δ*rsmA*. Here, we constructed a third strain as RsmA negatively regulates the translation of *rsmN* (21). When compared with RsmA::3×FLAG protein, RsmN::3×FLAG was not detected despite the deletion of translational repressor *rsmA* (Fig. S1A). Therefore, we expressed the *rsmN*::3×FLAG cassette from a multicopy plasmid in the PAO1 wild type strain. RsmN was successfully detected by western blot when the medium was supplemented with the arabinose inducer at 0.1% concentration (Fig. S1B). In the following procedure, chromosomally tagged *rsmA*::3×FLAG strain and plasmid-inducible *rsmN*::3×FLAG strain were incubated during the early stationary phase (OD_600_ = 2.0) and UV irradiated to induce covalent bonds between RNAs and the corresponding proteins *in vivo*. Autoradiography and western blot analyses indicated that UV crosslinking and co-immunoprecipitation with anti-FLAG antibodies along with stringent washing enriched RsmA-RNA and RsmN-RNA complexes (Fig. 1A). It should be noted that we omitted RNA radiolabeling steps and cut membranes from the regions consisting of RsmA/N bands up to the region 50 kDa above for RNA purification (Fig. 1B).

**Figure 1.**
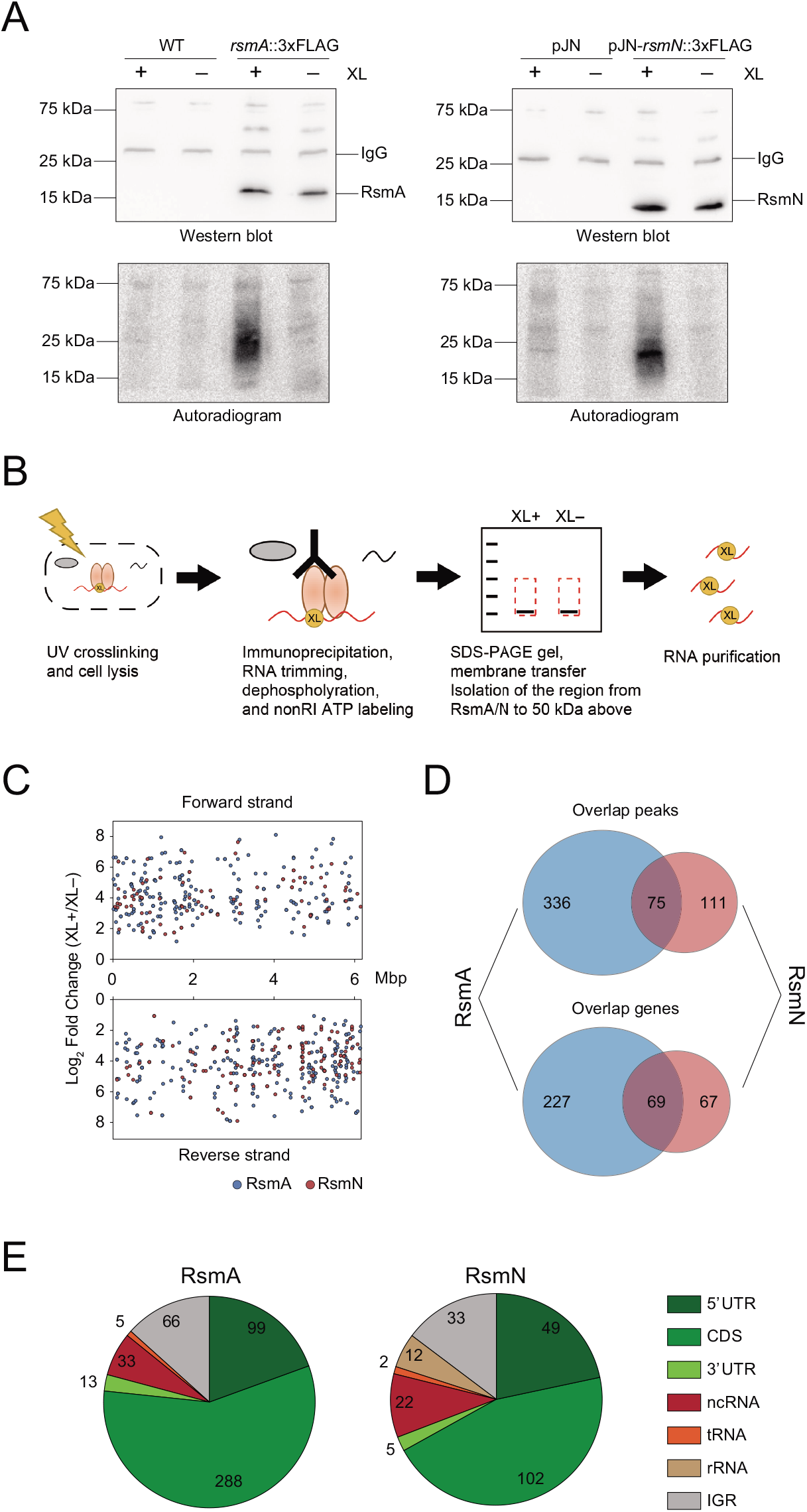
Overview of RsmA/N CLIP-seq. (A) Representative figures of autoradiogram and western blotting of the CLIP-enriched RsmA/N-RNA complexes. XL+: crosslinking, XL-: non-crosslinking. (B) Graphical summary of nonRI CLIP-seq approach. (C) The distribution of RsmA/N binding sites across the *P. aeruginosa* genome. Peaks from RsmA and RsmN CLIP-seq are highlighted in blue and red, respectively. (D) Overlapping peaks and genes between RsmA and RsmN bindings. Two peaks from both RNA chaperones wherein both the start and stop positions are within 40 nt were considered as the same peak. (E) Classification of RsmA/N peaks into RNA classes (5’ UTR, CDS, 3’ UTR, sRNA, tRNA, rRNA and Intergenic peaks). Note that a peak may be classified into multiple classes based on its position.

Purified RNAs from crosslinked and non-crosslinked samples in triplicates were reverse transcribed into cDNA and subjected to Illumina sequencing. By performing a comparative analysis of called peaks from RsmA/N-binding regions between crosslinked and non-crosslinked libraries using DESeq2, we detected hundreds of binding sites with significant enrichment (false discovery rate (FDR) < 0.05 and minimum expression = 1.0) throughout the *P. aeruginosa* transcriptome (Fig. 1C). We identified 75 overlapping peaks between RsmA and RsmN, with overlapping peaks defined as the peaks that exhibited both the start and stop positions within 40 nt of each other, thus constituting 69 overlapping genes (Fig. 1D). Since this CLIP-seq was carried out in conditions similar to the previously performed Hfq CLIP-seq (22), the genes that overlapped between RsmA/N and Hfq were also investigated. A total of 25 genes including the genes encoding outer membrane proteins were common among all three RNA chaperones (Fig. S2A). Gebhardt et al. recently performed chromatin immunoprecipitation with cells grown in the presence or absence of rifampicin followed by high-throughput DNA sequencing (ChIPPAR-seq) for *P. aeruginosa* RsmA to capture a subset of nascent transcripts (20). When compared with the list of transcripts that are regulated co-transcriptionally by RsmA, approximately 25% of RsmA-binding genes were also detected in our CLIP-seq (Fig. S2B). In addition, Romero et al. mapped more than 500 transcripts directly bound by RsmN using RNA immunoprecipitation and sequencing (RIP-seq) (19). We observed that a limited number of RsmN-binding genes were detected in both our CLIP-seq and the RIP-seq (Fig. S2C).

Finally, significant peaks were classified on the basis of RNA classes using previously generated UTR annotation list (22–24) and manually curated *P. aeruginosa* sRNA lists (25–27). Most of the peaks were mapped within mRNAs, in which more than 40% of the binding sites in CDSes overlapped with the first quarter of the CDSes (Fig. 1E and Fig. S3A). Interestingly, approximately half of ncRNAs bound by RsmA were annotated as *cis-*antisense RNAs (asRNAs) (Fig. S3B), suggesting that certain posttranscriptional regulatory mechanisms attributed to RsmA and asRNAs are still unrecognized. Taken together, RsmA/N CLIP-seq analysis identified hundreds of binding sites associated with each RNA chaperone.

### Similarities and differences in sequence and structural motifs bound by RsmA/N

RsmA/N bind to conserved GGA sequences within or close to Shine-Dalgarno sequences of target mRNAs (18). More precisely, parallel systematic evolution of ligands by exponential enrichment (SELEX) studies for RsmA/N have identified a common consensus motif CANGGAYG positioned in a hexaloop region of the stem-loop structure (28). To understand the consensus sequence bound by RsmA/N and its position in the target transcripts *in vivo*, the peak density of RsmA/N peaks across all detected mRNAs was determined via meta-gene analysis using start or stop codons as reference points. Strong peak densities were observed around start codons but not around stop codons, showing that *P. aeruginosa* RsmA/N preferentially bind to the 5’ UTRs of mRNAs (Fig. 2A). Next, sequence motifs for RsmA/N bindings were determined using all detected RsmA/N binding sequences by MEME motif analysis. Top-ranked motifs of both RsmA/N contained the ANGGA sequence (Fig. 2B). All RsmA binding sites exhibited the motif. In fact, more than 80% of detected RsmA/N peaks demonstrated at least one minimal GGA motif and the second nucleotide of ANGGA sequence for RsmA binding showed a preference for uracil (Fig. 2C). Unlike RsmA, the second nucleotide position of the ANGGA sequence for RsmN binding is more tolerant, accepting any nucleotides except for guanine (Fig. 2B and C). In order to understand whether the detected motifs were likely to be present in the apical loop, all detected RsmA/N binding sequences were subjected to CMfinder structural motif analysis. The top-ranked structural motifs from CMfinder were predicted as highly conserved stem-loop with the GGA sequence in the loop regions (Fig. 2D). Interestingly, the top-ranked structural motif for RsmN binding demonstrated two stem-loops with GGA sequences adjacent to each other, consistent with the previous report showing that RsmN requires two binding sites of the known target *tssA1* (28). Therefore, the number of GGA sequences per peak was searched to evaluate whether RsmN requires two adjacent GGA sequences for high-affinity binding. Although we expected that RsmN might tend to bind peaks corresponding to more than two GGA motifs, no significant difference was observed between RsmA and RsmN (Fig. 2E). Altogether, RsmA/N CLIP-seq demonstrates ANGGA in loop regions as a general recognition sequence/structural motif, which is in clear accordance with previously published works.

**Figure 2.**
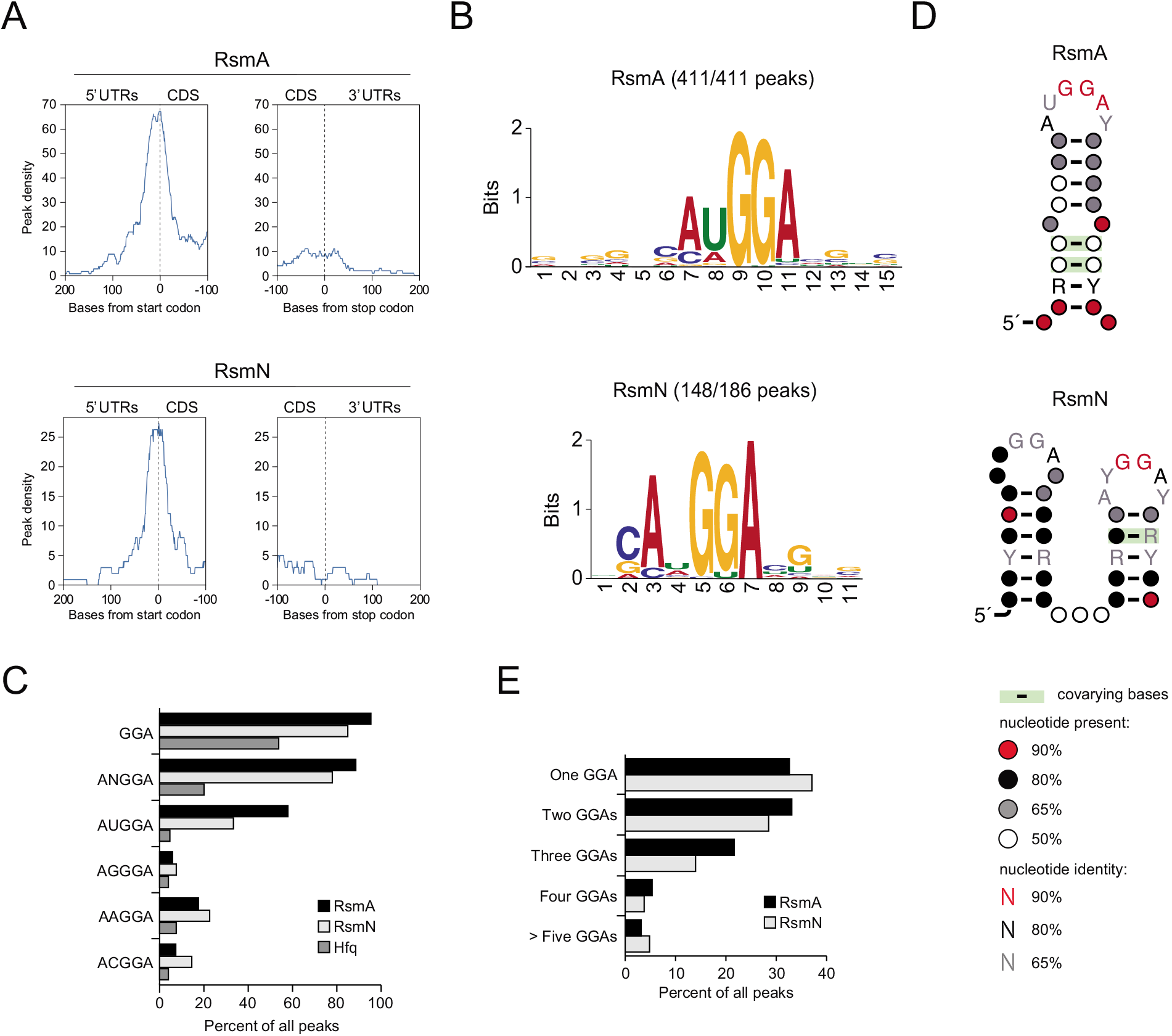
Consensus motif for *P. aeruginosa* RsmA/N bindings. (A) Meta-gene analysis for RsmA/N bindings along mRNAs with start and stop codons as the reference points. (B) MEME sequence motif analysis for all RsmA (411) and RsmN (186) peaks. The numbers indicate the peaks containing predicted sequence motifs. (C) Percentage of peaks with indicated sequences. (D) CMfinder structural motif analysis of all RsmA (411) and RsmN (186) peak sequences extended with 10 nt upstream and downstream. Top-ranked structural motifs are shown. (E) Percentage of peaks with the indicated number of GGA sequence per peak sequence extended with 10 nt upstream and downstream.

### RsmA/N CLIP-seq expands global binding targets in *P. aeruginosa*

When the small non-coding RNAs RsmY/Z are highly expressed, RsmA/N are titrated away from target RNAs playing an important role in the regulation of virulence factors associated with acute and chronic infections. The results of RsmA/N CLIP-seq conducted in this study demonstrate that the read coverage of RsmY/Z was highest among all other genes (Fig. 3A and Table S1). RsmY/Z exhibit multiple GGA motif sites for RsmA/N binding and RsmA/N peak sites corresponded to their GGA motifs (Fig. 3B), whereas Hfq predominantly associated with their Rho-independent terminators (29). Our data also shows that RsmA/N indeed bind to well-known target mRNAs. For examples, high CLIP-seq coverages were found at the 5’UTR of the gene *tssA1* encoding structural component of the Hcp secretion island-I-encoded T6SS and 5’UTR of the gene *mucA* encoding anti-sigma factor (Fig. 3C). The RsmA/N-binding sites detected on these mRNAs fold into hairpins with GGA motifs (Fig. 3D). Collectively, the data suggests that RsmA/N CLIP-seq was able to capture the *bona fide* RsmA/N binding sites.

**Figure 3.**
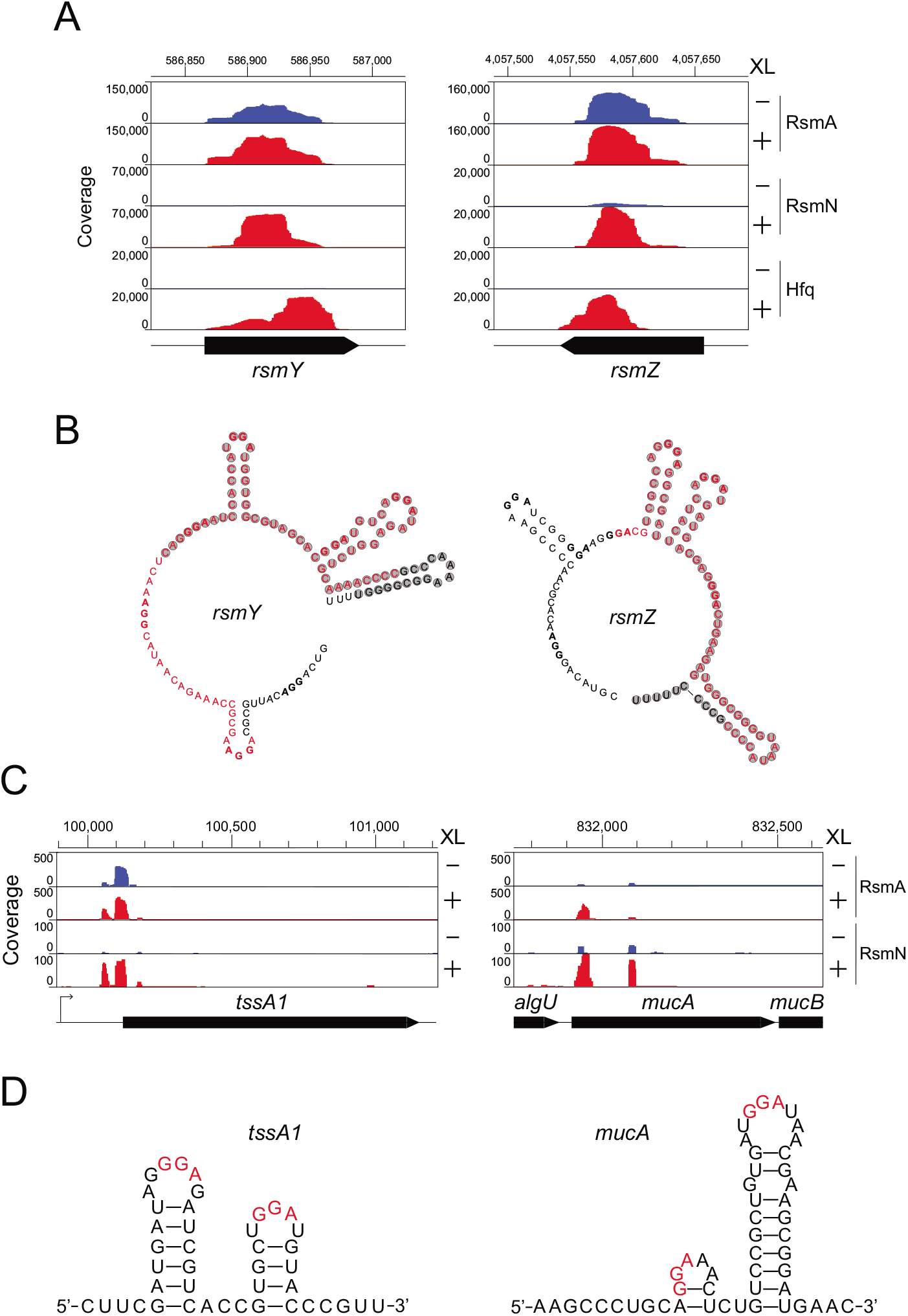
RsmA/N CLIP-seq captures previously known RsmA/N-binding sites. (A) Read coverage from RsmA/N and Hfq CLIP-seq at the sRNAs RsmY/Z loci. Vertical axis indicates each read count. XL+: crosslinking, XL-: non-crosslinking. (B) Secondary structures of sRNAs RsmY/Z. Red and shaded letters indicate RsmA/N and Hfq binding peaks, respectively. Bold letters indicate GGA sequence as a common binding motif of RsmA/N. (C) Read coverage at the *tssA1* and *mucA* loci from RsmA/N CLIP-seq are indicated. (D) Secondary structures of RsmA/N binding sites at the *tssA1* and *mucA* were predicted using Mfold (34). Red letters indicate GGA sequence as a common binding motif of RsmA/N.

To determine the biological processes in which RsmA/N-binding RNAs were enriched, DAVID enrichment analysis was performed for mRNAs containing peaks, with a modified Fisher’s exact *p*-value threshold < 0.1. Genes associated with cell wall organization, including those involved in the polysaccharide biosynthetic process, O-antigen biosynthetic process, and lipopolysaccharide (LPS) biosynthetic process were enriched (Fig. 4A). Among the cell wall organization processes, high peak density was detected within the *wbp* gene cluster involved in O5 O-antigen biosynthesis (Fig. 4B). Since the category represents an unknown role for RsmA/N, we decided to explore them in more detail.

**Figure 4.**
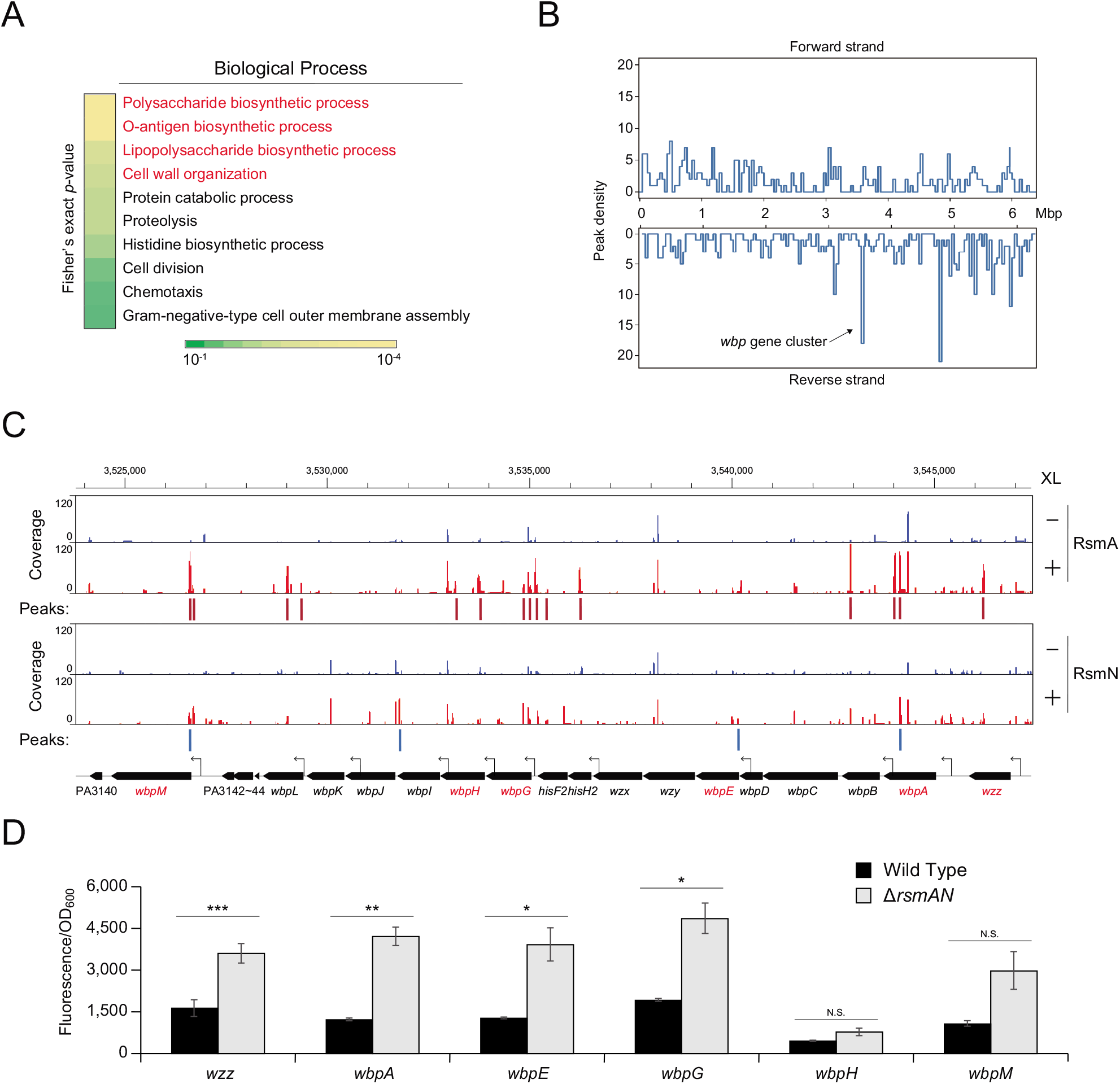
The *wbp* gene cluster is posttranscriptionally regulated by RsmA/N. (A) DAVID enrichment analysis of RsmA/N peaks. The results of biological process are presented. (B) RsmA/N peak density distribution along the *P. aeruginosa* chromosome in bins of 2 × 10^4^ basepairs. (C) Read coverage at the *wbp* gene cluster from RsmA/N CLIP-seq. TSS annotations (black arrows) were derived from a previous study (24). Vertical axis indicates each read count. RsmA and RsmN binding peaks with significant enrichment (FDR < 0.05) are indicated as red and blue bars, respectively. The genes shown in red were used for the following translational fusion assay. (D) Super-folder GFP translational fusion assay for the indicated genes between wild type and *rsmAN* mutant. Welch’s t-test results are indicated: *, *p*-value < 0.05; **, *p*-value < 0.01; ***, *p*-value < 0.001.

The *wbp* gene cluster consists of three genes responsible for the assembly of the O-antigen (*wzx, wzy*, and *wzz*), twelve genes involved in the biosynthesis assembly of the nucleotide sugars of the O unit (*wbpABCDEGHIJKLM*), three insertion genes encoding putative transposase and integrase (PA3142–PA3144) between *wbpL* and *wbpM*, and two non-LPS-related genes (*hisF2* and *hisH2*) between *wbpG* and *wzx* (30). Significant peaks indicating RsmA binding were observed in *wzz*, *wbpA, wbpB, hisH2, hisF2, wbpG, wbpH, wbpL*, and *wbpM* genes whereas those indicating RsmN binding were observed in *wbpA, wbpE, wbpI*, and *wbpM* genes (Fig. 4C). We verified the function with respect to RsmA/N-mediated gene regulation against *wzz*, *wbpA, wbpE, wbpG, wbpH* and *wbpM* using super-folder GFP (sfGFP) translational fusion assay. When compared with the wild type, Δ*rsmA/N* strain expressed significantly high GFP fluorescence in *wzz*, *wbpA*, *wbpE*, and *wbpG* translational fusions, suggesting that RsmA/N posttranscriptionally repress the translation of each gene (Fig. 4D). Altogether, our RsmA/N CLIP-seq results reveal new RsmA/N regulatory targets associated with LPS modification.

### Homeostasis of RsmY/Z expressions by an RsmA/N-controlled feedback loop

The expression of RsmA/N-titrating sRNAs RsmY/Z is activated by GacAS two-component system (15). The GacAS system is controlled by three additional HPKs: RetS, PA1611, and LadS. In addition to these complex regulatory pathways, RsmY and RsmZ are independently regulated by the histidine phosphotransfer protein HptB and cytoplasmic axial filament protein CafA, respectively. Expression of *hptB* is regulated by additional membrane-binding proteins SagS and ErcS (31), whereas the *cafA* expression is under the control of another two-component system BfiR/BfiS (32). Among those proteins involved in the homeostasis of RsmY/Z expression, we observed RsmA/N binding peaks in the *sagS*, *gacS, ladS, hptB*, and *cafA* genes (Fig. 5A). Since HptB is the decision hub for RsmY activation via the phosphorylation of additional regulatory components HsbA/HsbD and flagellar gene expression via FlgM sequestration (33), we focused on RsmA/N-mediated *hptB* gene regulation. We also investigated whether RsmA/N directly regulated *cafA* gene expression since CafA specifically represses RsmZ expression with its endoribonucleolytic activity.

**Figure 5.**
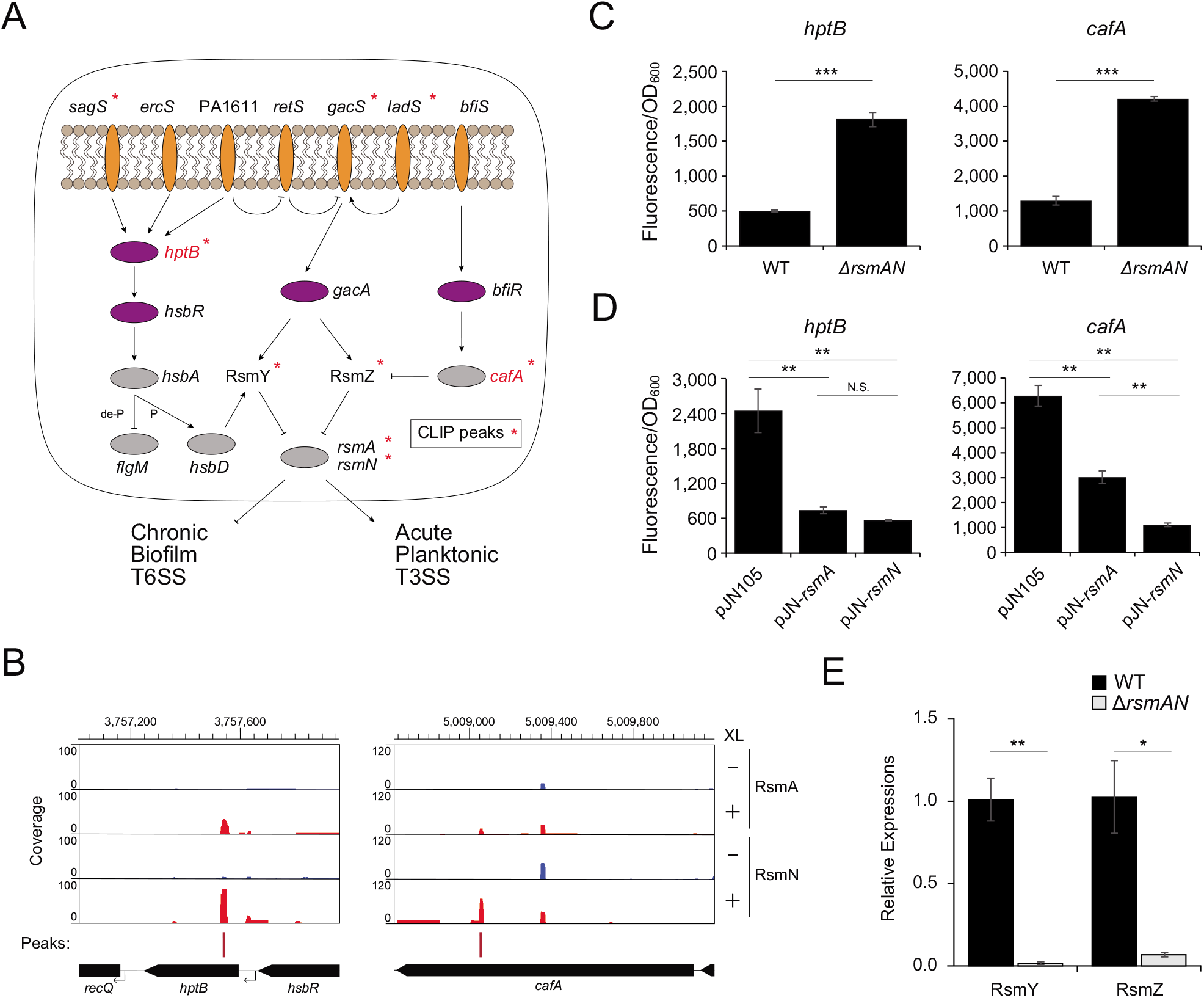
RsmA/N-controlled feedback loop for RsmY/Z sRNAs expressions. (A) Graphical summary of RsmA/N bindings with GacAS regulatory network. Orange and purple ovals are membrane-bound histidine kinases and response regulators in two-component systems, respectively. Asterisks indicate that RsmA/N peaks are associated with the genes. (B) Read coverage at the *cafA* and *hptB* loci from RsmA/N CLIP-seq. Vertical axis indicates each read count. RsmN binding peaks with significant enrichment (FDR < 0.05) are indicated as red bars. (C) Super-folder GFP translational fusion assay for the indicated genes between wild type and *rsmAN* mutant. (D) Super-folder GFP translational fusion assay for the indicated genes in *rsmAN* mutant with indicated plasmids. For exogenous RsmA/N expressions, 0.1% arabinose was added. (E) Relative expressions of RsmY/Z in PAO1 wild type and Δ*rsmA/N*. RNA was extracted from cultures at OD_600_ = 2.0 and RsmY/Z abundances were quantified by qRT-PCR. Welch’s t-test results are indicated: *, *p*-value < 0.05; **, *p*-value < 0.01; ***, *p*-value < 0.001 (C and E) and One-way ANOVA and Tukey’s HSD test results are indicated: **, *p*-value < 0.01; N.S., not significant (D).

We observed RsmA/N binding peaks in both coding DNA sequences although only RsmN binding was statistically significant (Fig. 5B). Therefore, we performed the sfGFP translational fusion assay to understand the RsmA/N associated gene regulation of *hptB* and *cafA*. When compared with the wild type, Δ*rsmA/N* strain expressed significantly high GFP fluorescence in both translational fusions (Fig. 5C). In order to investigate whether both RsmA and RsmN repress the *hptB* and *cafA* translations, RsmA and RsmN were exogenously expressed from multicopy plasmid pJN105 in Δ*rsmA/N* background and GFP fluorescence was measured. When compared with pJN105 vector control, both RsmA/N were capable of repressing GFP fluorescence (Fig. 5D). If *hptB* expression is negatively regulated by RsmA/N, downstream HptB-dependent secretion and biofilm anti anti-sigma factor HsbA should be phosphorylated and diguanylate cyclase HsbD should activate RsmY expression. Additionally, if *cafA* expression is negatively regulated by RsmA/N, degradation of RsmZ by CafA should be alleviated. To confirm these, RsmY/Z abundance between the wild type and Δ*rsmAN* strains was quantified by qRT-PCR. As expected, RsmY/Z were significantly reduced in the Δ*rsmAN* strain (Fig. 5E), suggesting that RsmY/Z expressions were balanced by RsmA/N-controlled feedback loop.

### RsmA/N bind to multiple sRNAs other than RsmY/Z in *P. aeruginosa*

Besides RsmY/Z, we observed high read coverage for RsmA/N bindings and significant enrichment of crosslinked samples in several sRNAs. For example, SPA0035 sRNA, a transcript from the minus strand of the intergenic region between *ada* and PA2119, was significantly bound to RsmA/N, but not to Hfq (Fig. 6A, left). As another example, the PAO1-specific asRNA SPA0066 transcribed from the opposite strand of PA3993 was also significantly bound to RsmA/N, but not to Hfq (Fig. 6A, middle). It should be noted that SPA0066 overlapped in the antisense orientation of the 5’ UTR of a gene encoding a putative transposase of the IS116/IS110/IS902 family. This transposase family exists in six genomic positions in PAO1, all of which demonstrate identical asRNAs (SPA0004–8 and SPA0066). The SPA0079 sRNA, a transcript from the minus strand of the intergenic region comprising 1,011 nt between PA2763 and PA2764, was significantly bound to RsmA/N, as well as to Hfq (Fig. 6A, right and Fig. S4A). The majority of these sRNAs carry GGA motifs and many corresponding peaks fold into hairpins (Fig. 6B).

**Figure 6.**
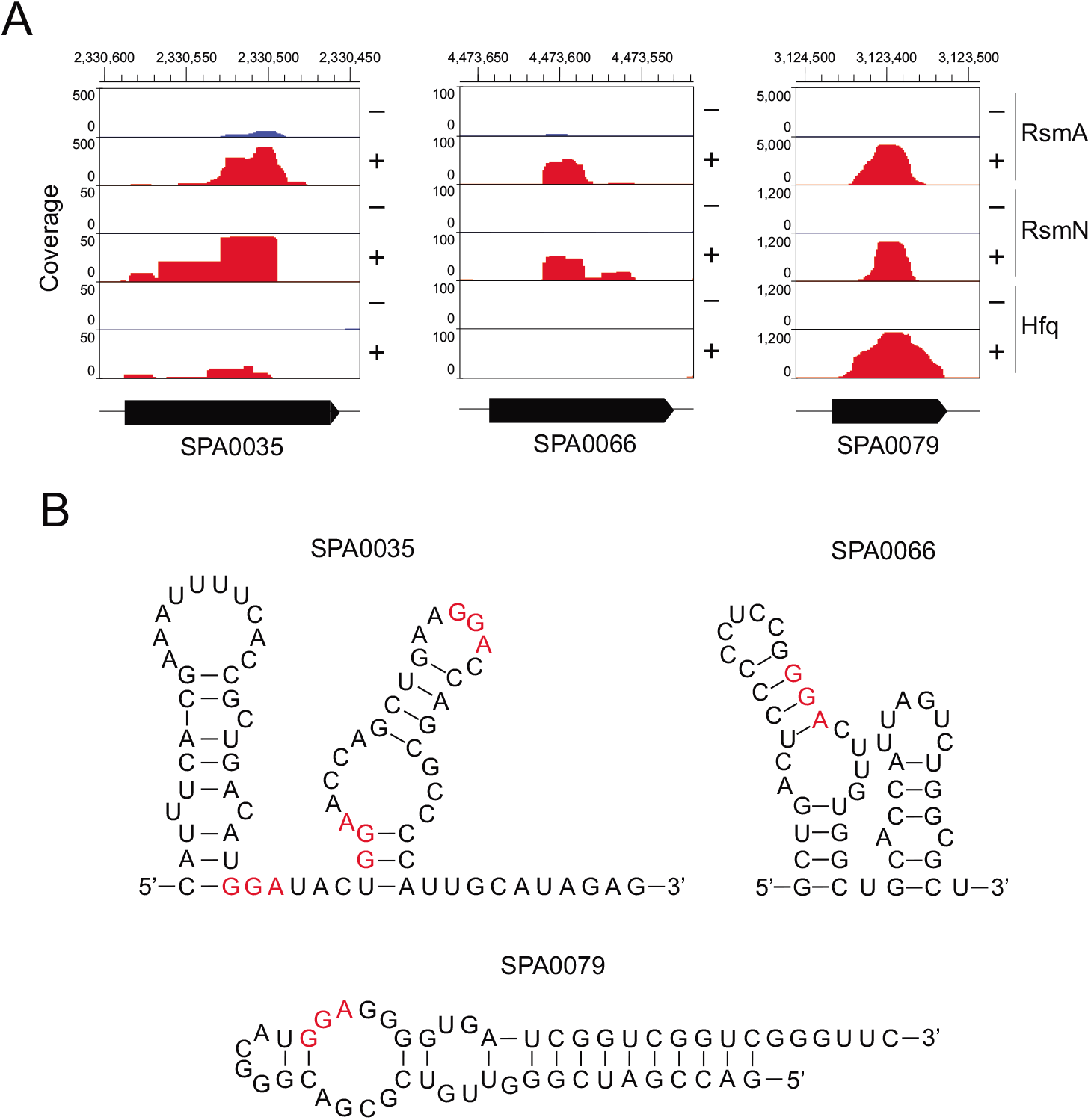
Representative RsmA/N-binding sRNAs. (A) Read coverage at the SPA0035, SPA0066, and SPA0079 loci from Hfq and RsmA/N CLIP-seq. (B) Secondary structures of binding sites of three sRNAs, which are extended by additional 10 nt upstream and downstream. Secondary structures were predicted using Mfold (34).

Among the RsmA/N-binding sRNAs detected in our CLIP-seq, SPA0079 exhibited significant binding to both Hfq and RsmA/N and demonstrated high coverage next to RsmY/Z (see y-axis in Fig. 6A and Table S1). Subsequently, we investigated its characteristics as an RsmA/N-binding sRNA in next section.

### SPA0079 sRNA does not sequester RsmA/N away from target mRNAs under our conditions

We first determined the SPA0079 boundaries by 5’/3’ RACE analysis. RNA pyrophosphohydrolase RppH treatment followed by PCR amplification selectively enriched 5’ terminus with triphosphate ends of SPA0079 (the band indicated by a red arrow in Fig. S4B). The 3’ terminus of SPA0079 corresponded to the canonical U repeats of the rho-independent terminator (Fig. S4A). The SPA0079 secondary structure predicted by Mfold (34) exhibited only a single stem-loop, having a strong resemblance to a rho-independent transcription terminator (Fig. S4C). The expression of SPA0079 in the wild type and the deletion mutant were measured throughout the course of growth (Fig. S4D). The maximal level of the SPA0079 expression was reached in the late stationary phase at the OD_600_ of 6.0 in the wild type (Fig. S4D, lanes 1–5). In contrast, SPA0079 sRNA level was undetectable in the ΔSPA0079 mutant strain (Fig. S4D, lane 6).

A previous study has suggested that two quorum sensing (QS) pathways LasI/LasR and RhlI/RhlR might be related to the expression of SPA0079 (35). Furthermore, the DNA sequence of the upstream of −35 promoter motif was similar to the consensus motif of LasR/RhlR binding (Fig. S4A). To investigate QS-dependent SPA0079 expression, northern blotting analysis was performed with Δ*lasl*Δ*rhlI* double mutant strain with or without the exogenous supplementation of two acyl homoserine lactones (AHLs) 3OC12-HSL and C4-HSL. SPA0079 displayed strict dependence on QS as it accumulated within 1 h after AHLs were added (Fig. S5A). Additionally, SPA0079 was not detectable in a Δ*lasI* strain, whereas it was detected moderately in a Δ*rhlI* strain (Fig. S5B). Taken together with the published report (35), these results suggest that the expression of SPA0079 is controlled by the QS systems. Another previous report showed that Hfq stabilizes RsmY by blocking the cleavage by RNase E (29). We investigated whether Hfq might affect SPA0079. Northern blotting showed no difference in the SPA0079 levels in a Δ*hfq* mutant compared to its levels in wild type and Δ*rsmA/N* mutant (Fig. S5C).

To investigate whether SPA0079 indeed titrates RsmA/N away from the mRNA target, the activity of sfGFP fused with 5’ UTR and CDS of *cafA* was assayed with or without SPA0079 overexpression. Although we speculated that a *cafA*::sfGFP fusion would be repressed by the induction of SPA0079, no clear difference was observed in GFP activity in the SPA0079 overexpression state compared to that in the Δ*rsmA/N* mutant strain (Fig. S5D). Considering that the level of SPA0079 expression reaches maximum in the late stationary phase (Fig. S4D), and that RsmY/Z predominantly bind to RsmA/N under the conditions in which CLIP-seq was performed, it may be that SPA0079 sRNA competes or complements RsmY/Z under specific conditions not captured in our assay. Future studies are required to determine whether or not SPA0079 is a functional RsmA/N antagonist.

## DISCUSSION

Two conserved and redundant RBPs RsmA/N play a critical role in balancing acute and chronic infections in *P. aeruginosa*. So far, the interactions of RsmA/N with target RNAs have been extensively explored *in vitro*. However, it is prudent to understand the mode of actions in the complex transcriptional networks *in vivo*. Many studies using advanced technologies based on high-throughput sequencing have now provided the global regulatory functions of RBPs depending on the physiological states of a cell (36–38). In this study, we performed the RsmA/N CLIP-seq analysis and demonstrated more than 500 genome-wide RsmA/N binding sites. Many genes identified in this study could be attractive targets for further elucidating the regulatory mechanisms of RsmA/N in *P. aeruginosa*.

As the prerequisite for this study, we first checked RsmA and RsmN abundance at an OD_600_ = 2.0. It is well known that the transcript encoding RsmN is posttranscriptionally repressed by RsmA and the protein abundance is relatively low compared to RsmA during the growth period (21). Therefore, we speculated that RsmN-binding RNAs are likely captured in the Δ*rsmA* background. However, the abundance of RsmN is still low even in the Δ*rsmA* background and interacting RNAs cannot be captured by co-immunoprecipitation (Fig. S1). The low RsmN abundance may be attributed to its original low transcription rate in the condition where our CLIP-seq was conducted. A previous study has shown that the level of *rsmN* expression reaches maximum in the late stationary phase (39). Although we relied on the exogenous expression of RsmN from multicopy plasmid and performed RsmA/N CLIP-seq at an OD_600_ = 2.0 in order to compare previously performed Hfq CLIP-seq result (22), it would be possible to detect RsmN-binding positions using the chromosomally tagged strain at the later growth phase, which should be addressed in further experiments.

In the widespread application of the current CLIP-seq procedure, radioactive labeling of crosslinked RNAs and visualization of the autoradiograph are common. As with several non-radioactive CLIP methods (40–43), herein, we omitted labor-intensive use of radioactive substances and successfully captured specifically interacting RNAs (Fig. 1B to E). When compared with previously published data, RsmA/N-binding genes demonstrated in this study were relatively low in number (Fig. S2). This variation may be attributed to the differences in the experimental setup. RsmA ChIPPAR-seq captures nascent transcript at mid-log phase and significant binding positions are defined as enrichment in the minus rifampicin library compared with the plus rifampicin library (20). RsmN RIP-seq is performed during the late stationary phase and significant binding genes are defined using enrichment index generated by comparison of co-immunoprecipitation library with total RNA library (19). In addition, the small overlap of detected genes with previous reports perhaps derives from the limitations of bioinformatics analysis. Our CLIP-seq approach relies on the comparison between crosslinking and non-crosslinking libraries. Hence, this approach identifies the binding sites with significant enrichment in crosslinking samples. The analysis perhaps missed the *bona fide* RsmA/N binding sites with high-affinity in non-crosslinking libraries. Especially in our RsmA CLIP-seq, the background likely has much higher counts than that specific to the crosslinking libraries, which might be the artifact of one or two highly expressed transcripts such as RsmY and RsmZ (Fig. 3A and Fig. S6). Nevertheless, numerous known targets of RsmA/N were observed from our CLIP-seq (Fig. 1). As examples, we showed high CLIP-seq coverages observed at the 5’UTRs of the mRNAs *tssA1* and *mucA* and sRNAs RsmY/Z, leading to the identification of their RsmA/N-binding sites *in vivo* at a single-nucleotide resolution (Fig. 3).

Interestingly, approximately half of RsmA-binding ncRNAs were annotated as asRNAs (Fig. S3B). Although no report exists demonstrating the promotion of the complex formation between regulatory RNAs and their target mRNAs in Gram-negative bacteria by the CsrA family, *Bacillus subtilis* CsrA helps SR1 sRNA basepair with *ahrC* mRNA, affecting the expression of its downstream genes (44). Rather than the CsrA family, ProQ/FinO-domain proteins and Hfq are generally thought to be responsible for the interaction between mRNAs and asRNAs in Gram-negative bacteria (45). The *E. coli* F plasmid-encoded FinO promotes the association of F plasmid-encoded mRNA *traJ* with asRNAs FinP (46). The chromosomally encoded ProQ from *Salmonella* also promotes the association of mRNA with asRNAs including Type I Toxin-Antitoxin system (47, 48), although ProQ can also play a similar role in Hfq acting as a matchmaker between *trans*-encoded sRNAs and mRNAs (49). Hfq primarily promotes the baseparing between *trans*-encoded sRNAs and mRNAs. However, it also facilitates antisense paring such as in IS10 system, where asRNA RNA-OUT interacts with RNA-IN encoding transposase to occlude the ribosome binding site (50). Since we found that asRNAs SPA0004–0008 and SPA0066 transcribed from the opposite strands of genes encoding putative transposases associate with RsmA/N (Fig. 6A, middle), it should be interesting to investigate whether these asRNAs may regulate the opposite genes via RsmA/N binding similar to Hfq, RNA-IN, and RNA-OUT in *E. coli*.

The advantage of CLIP-seq is to detect protein binding sites on the transcript at a single-nucleotide resolution. This approach has demonstrated sequence/structural binding motifs of the CsrA family in *E. coli* and *Salmonella* (37, 38). Herein, we investigated the similarities and differences between *P. aeruginosa* RsmA and RsmN on the basis of sequence/structural motifs and a typical distribution of the detected peaks (Fig. 2). The results of the peak distribution among mRNAs were common between the two RBPs, indicating that RsmA/N peaks are highly enriched at only 5’ UTRs, which is consistent with the previous reports (19, 51) (Fig. 2A to D). In addition, sequence and structural motif analyses demonstrated that the stem-loop structure with the ANGGA sequence in its loop region is primary of RsmA/N bindings. Interestingly, although all of the RsmA peaks detected in this study include the ANGGA motif, RsmN peaks seem to be under looser definition of its binding motif than RsmA (see the number of detected peaks in Fig. 2B). From the perspective of molecular basis, RsmA homolog RsmE in *Pseudomonas fluorescens* CHA0 needs the looped out nucleotide base N in ANGGA motif and baseparing in the stem for rigid binding through Leu55 to Ala57 in the C-terminal α helix as well as the correct stacking (52). In contrast, α helix in RsmN is located within the internalized β2 and β3 sheets, constituting the hydrophobic core that acts as a potential new RNA binding site (13). Thus, RsmN may bind with a more flexible loop motif independent of C-terminal α helix even without obvious ANGGA sequence. Interestingly, the structural motif of RsmN peaks consisted of two tandem stem-loop structures with GGA motifs (Fig. 2D). Consistent with this, *in vitro* SELEX study and mutational analysis have demonstrated long RsmA/N targets with two consensus GGA motifs, and both of the two tandem GGA sites are required only for RsmN binding (28). Overall, our CLIP-seq analysis combined with previous reports suggested a unique two-sidedness of RsmN binding; tandem stem-loop structures with two GGAs or flexible sequences without obvious ANGGA sequence.

Further genetic analysis has elucidated new regulatory targets of RsmA/N involved in O5 O-antigen biosynthesis (Fig. 4). The three genes *wzx*, *wzy*, and *wzz* are responsible for the assembly of the O-antigen and other genes *wbpABCDEGHIJKLM* are involved in the biosynthesis assembly of the nucleotide sugars of the O unit (30). Since we observed RsmA bindings to both *wzz* and *wzz2* encoding O-antigen chain length regulators and validated the RsmA/N-mediated translational repression of *wzz* (Fig. 4D and Table S1), active posttranscriptional regulation through RsmA/N would lead to LPS-rough phenotype, which is often observed in *P. aeruginosa* isolates from chronic pulmonary infections (53, 54). In a recent study, the low levels of *wzz2* expression led to the conversion to mucoid phenotype with alginate production, and a transcriptional factor AmrZ negatively regulated the expression (55). Under the same AlgU regulon with AmrZ, AlgR activates alginate production and in turn, directly stimulates RsmA expression (56). This might accelerate the modulation of O-antigen length and eventually the biofilm formation (57). The translational repression of genes involved in O-antigen biosynthesis through RsmA/N is also consistent with the findings demonstrating that the absence of B-band O-antigen is correlated with the increase of T3SS-mediated cytotoxicity that is activated by RsmA/N directly (58). Taking into account the previous ChIP-seq analysis showing that T3SS master regulator ExsA binds to promoter sequences of *wbpA* and *wbpH* (59), the *wbp* gene cluster is likely to be under the both transcriptional and translational regulation mediated by RsmA/N.

Free RsmA/N and the level of the titrating sRNAs are controlled through homeostatic regulation. Although it was previously shown that RsmA exerts a positive effect on RsmY and RsmZ transcription in *P. aeruginosa* and *Pseudomonas protegens* (60, 61), the mechanisms underlying the positive feedback loop remain unclear. In this study, *hptB* encoding the histidine phosphotransfer protein and *cafA* encoding the cytoplasmic axial filament protein were identified as novel RsmA/N regulatory targets (Fig. 5). When RsmA/N repress the translation of *hptB*, RsmY expression would be stimulated via the phosphorylation of HsbA and the subsequent activation of HsbD (31). In addition to RsmY, when the *cafA* is repressed in the RsmA/N-dependent manner, RsmZ would escape from degradation owing to the endoribonucleolytic activity (32), thus constituting three repressors feedback loop. In a simple and synthetic gene circuit, however, the three repressors loop periodically induces the synthesis of output (62). Since RsmY/Z expression increases throughout the growth period (15), other factors might be involved in the homeostatic regulations of RsmY/Z. For example, polynucleotide phosphorylase PNPase regulates the stabilities of RsmY/Z (63). Our CLIP-seq data has identified RsmA binding to the gene encoding PNPase (Table S1), which infers that RsmA affects RsmY/Z through the availability of PNPase.

Finally, we discovered 41 RsmA/N-binding sRNAs besides known RsmA/N-titrating sRNAs (Fig. 6 and Table S1). Among them, SPA0079 was highly enriched in both Hfq and RsmA/N CLIP-seq, motivating us to further investigate its properties. The level of SPA0079 expression reached maximum in late stationary phase and was undetectable in Δ*lasIΔrhlI* mutant, whereas exogenous supplementation of two AHLs complimented the expression (Fig. S4 and S5), strongly supporting the fact that SPA0079 is regulated by two QS pathways (35). Although we expected that SPA0079 could alleviate RsmA/N-mediated translational repression, no clear activation of translational fusion activity was observed with the SPA0079 sRNA overexpression (Fig. S5D). Given that RsmY/Z predominantly bind to RsmA/N under the conditions in which CLIP-seq was performed and the effect of SPA0079 is negligible, future studies should be performed using Δ*rsmY/Z* mutant to determine whether or not the SPA0079 sRNA are indeed RsmA/N antagonist. A recent paper shows that sRNA179 (also annotated as SPA0034) expression stimulates RsmY transcription (64). Additionally, the expression of sRNA179 is activated by QS similar to SPA0079 (35). Considering these observations, QS might suppress the RsmA/N regulatory system through two different posttranscriptional pathways, highlighting the complexity of Rsm regulatory systems.

## METHODS

### Strains, plasmids, and growth conditions

Strains, plasmids and oligonucleotides used herein are enlisted in Table S2. All experiments were performed using *P. aeruginosa* PAO1 or its derived strains. Each strain was cultured at 37°C in Luria-Bertani (LB) medium. Samples were collected at the OD_600_ values indicated in the figures. Where indicated, the appropriate AHLs were added at the following final concentrations: C4-HSL (Cayman Chemicals) at 10 μM and 3OC_12_-HSL (RTI International) at 2 μM. Antibiotics and arabinose were used as needed at concentrations listed as follows. For *Pseudomonas*, gentamicin, tetracycline, and arabinose at concentrations 50 μg/ml, 80 μg/ml, and 0.1%, respectively, were used. Similarly, for *E. coli*, gentamicin and tetracycline at concentrations 10 μg/ml and 20 μg/ml, respectively, were used.

### Strain construction

*P. aeruginosa* PAO1 Δ*rsmA*, Δ*rsmN*, Δ*rsmA/N* double mutant, Δ*lasI*, Δ*rhlI*, Δ*lasI*Δ*rhlI* double mutant, and ΔSPA0079 strains were constructed based on the conjugative transfer of appropriate plasmids and homologous recombination between chromosome and plasmids as described previously (65). The plasmids for the homologous recombination of Δ*rsmA*, Δ*rsmN*, and Δ*rsmA/N* double mutant were constructed by HindIII/XbaI cloning of PCR products 500 bp upstream and 500 bp downstream of CDS of each gene from PAO1 chromosome into pG19II backbone (66). The plasmids for the homologous recombination of Δ*lasI*, Δ*rhlI*, Δ*lasI*Δ*rhlI* double mutant, and ΔSPA0079 were constructed by HindIII/BamHI cloning of PCR products 500 bp upstream and 500 bp downstream of CDS of each gene from PAO1 chromosome into pG19II backbone (66). pG19*rsmA*::3×FLAG and pG19*rsmN*::3×FLAG were constructed by cloning PCR products at HindIII/XbaI sites 500 bp upstream of the *rsmA/N* stop codon in the PAO1 chromosome, 3×FLAG tag, 500 bp downstream of the *rsmA/N* stop codon in the PAO1 chromosome into the pG19II backbone.

### UV crosslinking, immunoprecipitation, and RNA purification

Bacterial cultures with volume 200 ml of three replicates were maintained up to an OD_600_ of 2.0 in LB medium. For RsmN expression from the multicopy plasmid, 0.1% arabinose was added at an OD_600_ of 0.8. UV crosslinking and immunoprecipitation were performed in accordance with the previously published protocol with minor modifications (22). Briefly, half of the culture at an OD_600_ of 2.0 from each condition was irradiated at 800 mJ of UV light at 254 nm. After UV crosslinking, the sample was centrifuged at 3,600 rpm for 30 min at 4°C along with the non-crosslinked control samples.

Cell pellets were lysed in FastPrep 24 (MP-Biomedicals) at 6 m/s for 1 min with 1 ml of 0.1-mm glass beads and 800 μl NP-T buffer (50 mM NaH_2_PO_4_, 300 mM NaCl, 0.05% Tween20, pH 8.0). NP-T supplemented with 8 M urea was added to each supernatant at an equal volume and incubated for 5 min at 65°C with shaking at 900 rpm. Anti-FLAG magnetic beads were washed thrice with 500 μl NP-T buffer, resuspended in 125 μl NP-T buffer, and treated with a 120-μl suspension of urea-treated samples. After 1 h of incubation at 4°C, samples were washed twice with 500 μl high-salt buffer (50 mM NaH_2_PO_4_, 1M NaCl, 0.05% Tween20, pH 8.0), followed by two washes with 500 μl NP-T buffer. Beads were resuspended in benzonase mix [500 units benzonase nuclease (E1014, Sigma-Aldrich) in NP-T buffer with 1 mM MgCl2] and incubated for 10 min at 37°C with shaking at 900 rpm. After one wash with high-salt buffer and two washes with CIP buffer (100 mM NaCl, 50 mM Tris-HCl, pH 7.4, 10 mM MgCl2), beads were resuspended in 200 μl CIP mix (20 units of calf intestinal alkaline phosphatase (M0290, NEB) in CIP buffer) and incubated for 30 min at 37°C with shaking at 800 rpm.

Here, radioactive isotope (RI) labeling was performed only in the preliminary investigation to check whether UV crosslinking did enrich RNA-protein complex (Fig. 1A). After one wash with high-salt buffer and two washes with PNK buffer (50 mM Tris-HCl, pH 7.4, 10 mM MgCl2, 0.1 mM spermidine), beads were resuspended in 200 μl PNK buffer and separated into 20 μl and 180 μl volumes to be used subsequently for western blotting and RI labeling, respectively. For RI labeling, beads were magnetized and resuspended in 100 μl PNK mix [10 units of T4 polynucleotide kinase (EK0032, ThermoFisher Scientific), 10 μCi γ-^32^P-ATP in PNK buffer]) and incubated for 30 min at 37°C, following the addition of 1 μl 10 mM non-radioactive ATP and incubation for 5 min at 37°C. After two washes with NP-T buffer, beads were resuspended in 25 μl 2xProtein loading buffer and incubated for 5 min at 95°C. Beads were magnetized and supernatants were transferred to fresh tubes. The elution was repeated twice. For western blotting, 20 μl separated beads were washed twice with 100 μl NP-T buffer and resuspended in 10 μl 2xProtein loading buffer and incubated for 5 min at 95°C. Beads were magnetized and supernatants were transferred to fresh tubes. The elution was repeated twice.

In the CLIP experiments for RNA purification and cDNA preparation, RI labeling was not used (Fig. 1B). After CIP reaction, beads were resuspended in 100 μl PNK mix [10 units of T4 polynucleotide kinase (EK0032, ThermoFisher Scientific), 1μl 10 mM non-radioactive ATP in PNK buffer] and incubated for 30 min at 37°C, followed by addition of 1 μl 10 mM non-radioactive ATP and incubation for 5 min at 37°C. After two washes with NP-T buffer, beads were resuspended in 30 μl 2xProtein loading buffer and incubated for 5 min at 95°C. Beads were magnetized and supernatants were transferred to fresh tubes. The elution was repeated twice.

Aliquots of 55 μl were loaded and separated via SDS-polyacrylamide gel electrophoresis (12% resolving gel) at 30 mA while moving through the stacking gel, which was then increased to 40 mA. After RNA electrophoresis, the RNA was transferred from the gel to the Protran membrane (#10600016, GE Healthcare). The membrane was placed on a clean glass surface and cut from the prestained protein markers to 50 kDa above them. Each membrane piece was cut into smaller pieces and placed in 2 ml fresh tubes with 400 μl PK solution [1.3 mg/ml Proteinase K (EO0491, ThermoFisher Scientific), and 10 units of RNase inhibitor (#10777019, Invitrogen) in 2xPK buffer (100 mM Tris-HCl, pH 7.9, 10 mM EDTA, 1% SDS)], followed by incubation for 1 h at 37°C with shaking at 1,000 rpm. The samples were subsequently incubated with 100 μl PK buffer containing 9 M urea for 1 h at 37°C with shaking at 1,000 rpm. Thereafter, 450 μl of supernatants from proteinase K-treated membranes were mixed with an equal volume of phenol:chloroform:isoamyl alcohol in phase-lock gel tubes and incubated for 5 min at 30°C with shaking at 1,000 rpm. Each mixture was centrifuged for 12 min at 13,000 rpm at 4°C, and 400 μl of the aqueous phase was precipitated with 3 volumes of ice-cold ethanol, 1/30 volume of 3 M NaOAc (pH 5.2), and 1 μl of GlycoBlue (AM9515, Invitrogen) for 2 h at −20°C. The precipitated pellet was washed with 80% ethanol, briefly dried for 5 min at 20°C, and resuspended in 11 μl of sterilized water.

### cDNA library preparation and sequencing

A cDNA library of the CLIP samples was prepared using the NEBNext Multiplex Small RNA Library Prep Set for Illumina (#E7300, NEB) in accordance with the manufacturer’s instructions. RT primer and both 3’/5’ SR adapters were diluted 10-fold with nuclease-free water before use. cDNA was converted from 2.5 μl of each purified RNA. Reverse-transcribed cDNAs were amplified by 20 cycles PCR. PCR products were concentrated to 10 μl using the MinElute PCR Purification kit and separated by 6% polyacrylamide gel with 7 M urea. Bands between 130 bp and 250 bp were cut and purified from the gel in accordance with the manufacturer’s instructions. Purified cDNAs of 5 μl volume were amplified by 6 cycles PCR and concentrated to 10 μl using the MinElute PCR Purification kit. Final cDNA libraries were quantified using the Qubit dsDNA assay (Q32854, Invitrogen) and Agilent 2100 Bioanalyzer DNA HS (Agilent). Twelve cDNA libraries from CLIP were pooled on an Illumina HiSeq2500 and sequenced in paired-end mode (2 × 50 cycles).

### Sequence processing, mapping, normalization, and peak calling

To assure high sequence quality, read 1 (R1) and read 2 (R2) files containing the Illumina paired-end reads in FASTQ format were quality and adapter-trimmed by Cutadapt (67) version 1.15/1.16 using a cut-off Phred score of 20. Reads without any remaining bases were discarded (command line parameters: −q 20 −m 1 −a AGATCGGAAGAGCACACGTCTGAACTCCAGTCAC-A GATCGTCGGACTGTAGAACTCTGAACGTGTAGATCTCGGTGGTCGCCGTATCATT). To eliminate putative PCR duplicates, paired-end reads were collapsed using FastUniq (68) After trimming, we applied the pipeline READemption (69) version 0.4.5 to align all reads longer than 11 nt to the *P. aeruginosa* PAO1 chromosome (NCBI accession no. NC_002516.2) reference genome using segemehl (70) version 0.2.0 with accuracy cut-off of 80%.

Read counts per position were analyzed using reads that mapped uniquely to single genomic position. The core of positions present in crosslinked and non-crosslinked library pairs were isolated, as described previously (71). Positions with low read counts were filtered from both crosslinked and non-crosslinked libraries using 10 standard deviations from 0 as a threshold. After plotting the difference in read counts between the two libraries as shown in Fig. S6, the size factor was calculated using the DESeq normalization procedure from the high-count positions in both libraries across all replicates (72).

We applied PEAKachu v0.1.0 (https://github.com/tbischler/PEAKachu) for the peak calling similar to the previously described protocol (71). First, BAM files for the respective pairs of crosslinked and non-crosslinked libraries were used to run in paired-end (-P) and paired-replicates (-r) mode. The maximum fragment size (-M) was set to 50 and annotations generated as GFF format were used to map overlapping features to the called peaks. Normalization was performed in the ‘‘manual’’ mode using previously determined size factors (see above). Other parameters were set to default. Second, the boundary of initial peaks was set through block definition computed by the blockbuster algorithm (73) based on pooled read alignments from all crosslinked libraries using default parameters. Third, the PEAKachu tool ran the DESeq2 package (74) to analyze the significance of peak enrichment in the crosslinked libraries relative to the non-crosslinked libraries with parameter values as follows: mad-multiplier (-m) 1.0, fold change (-f) 1.0, and adjusted *p*-value (-Q) 0.05. Finally, PEAKachu was used for each replicon and strand to generate normalized coverage plots to facilitate data visualization.

### Analysis of sequence and structural motifs

The sequences of peaks were used to perform MEME sequence motif analysis (75). Minimum and maximum motif widths were set at 6 and 50, respectively, while other parameters were set to default. Structural motifs of the sequences of peak regions extended by an additional 10 nt upstream and downstream were analyzed using CMfinder 0.2.1 (76). The minimum length of single stem-loop candidates was set to 20, while other parameters were set to default. Each analyzed motif was visualized using R2R (77).

### Statistical and other analysis

Descriptive statistical analyses for peak and gene overlapping between RsmA and RsmN, peak distributions across the *P. aeruginosa* genome, and peak classification among RNA classes were performed using Microsoft Excel. Meta-gene analyses across detected mRNAs or the *P. aeruginosa* genome were performed with in-house script using Python3. Genes identified via CLIP-seq analysis were functionally characterized using the PseudoCAP annotation (http://www.pseudomonas.com). KEGG enrichment analysis was performed for mRNAs identified via CLIP-seq analysis using DAVID (https://david.ncifcrf.gov/summary.jsp). Default parameters and databases were used for the analysis.

### Construction of translational fusion plasmids

The sequence containing the first 186 amino acids of LacZ (LacZ186) and the super-folder GFP (sfGFP) open reading frame (78) expressed by J23119 constitutive promoter was obtained from ThermoFisher Scientific. LacZ186 was flanked by a NsilI and a SphI site while sfGFP was flanked by a SphI and an XbaI site. The synthetic sequence was cloned into inversely amplified pSW002-Pc PCR product by infusion cloning (Takara, Z9633N). pSW002-Pc was a gift from Rosemarie Wilton (Addgene plasmid # 108234) (79). The region from TSSes to downstream of the start codon in target RNAs was amplified with 15 bp complemented sequences on 5’ and 3’ end, respectively. The resulting PCR products were cloned into a NsilI/SphI digested pSW-*lacZ*::sfGFP backbone by infusion cloning (Takara, Z9633N).

### Translational fusion assay

*P. aeruginosa* wild type and Δ*rsmA/N* strains carrying the sfGFP translational fusions were grown overnight in 1 ml LB medium with 80 μg/ml tetracycline at 37°C. An aliquot with 2 μl volume was resuspended in 200 μl LB medium with 80 μg/ml tetracycline and transferred to black polystyrene 96-well microplates with a clear, flat bottom (Corning). The medium was additionally supplemented with gentamicin with concentration 50 μg/ml and 0.1% arabinose for RsmA/N and SPA0079 expression from multicopy plasmids. Fluorescence polarization (FP^476/510^) was measured with SpectraMax GeminiXS (Molecular Devices) every 15 min for 6 h at 37°C with agitation for 1 min before fluorescence measurement. The fluorescent cutoff was set to 495 nm. Thereafter, the final optical density (OD) of each culture was obtained by GeneQuant 1300 (Biochrom). GFP activity was expressed in arbitrary units as FP_476/510_/OD_600_.

### qRT-PCR

Total RNA was extracted by the hot phenol extraction method followed by DNase treatment. cDNAs from 1 μg DNase-treated purified RNA were obtained using PrimeScriptTM RT Master Mix (#RR036A, TAKARA) following the manufacturer’s instruction. qRT-PCR was performed for triplicates. Reactions were performed in a total volume of 20 μl containing 10 μl of TB Green Fast qPCR Mix (#RR430A, TAKARA), 0.4 μl of ROX reference dye II, 0.8 μl of each primer (10 μM), 7 μl of RNase-free water and 1 μl of template DNA. PCR and fluorescence measurements were performed using the viiA7 (Applied Biosystems) with the following program; preheating at 95°C for 30 s, followed by 40 cycles of denaturation at 95°C for 5 s and annealing at 60°C for 20 s. The gene *rpoD* was used as an internal standard (80). Relative gene expression was calculated using the ΔΔCt method (81).

### 5’ and 3’ rapid amplification of cDNA ends (RACE)

5’ and 3’ RACE assays were performed as previously described with some modifications (82). For 5’ RACE assay, 15 μg of DNA-depleted RNA was incubated with 12.5 units of RNA 5’ pyrophosphohydrolase (RppH) (#M0356, NEB) in a 50 μl reaction for 1 h at 37°C. The same volume of RNA without the RppH reaction was prepared to be used as a negative control. After incubation, RppH-reacted and control RNAs were purified and 1.25 μM 5’RACE adaptor (5’-GAU AUG CGC GAA UUC CUG UAG AAC GAA CAC UAG AAG AAA-3’) was mixed. The samples were denatured at 95°C for 5 min. The adaptor was ligated using T4 RNA Ligase 1 overnight at 16°C. The adaptor-ligated RNA was purified, annealed with gene-specific primer and reverse-transcribed using AffinityScript Multiple Temperature Reverse Transcriptase (#600107, Agilent) with following conditions; 42°C for 20 min, 55°C for 20 min and 70°C for 15 min in a 20 μl reaction. One-micro-liter of reverse-transcribed cDNA was amplified by nested PCR with a total volume of 50 μl containing 0.4 μM of 5’adaptor-specific primer, 0.4 μM of 1^st^ round gene-specific primer, 200 μM of each dNTP, 1x PCR buffer, and 1.25 units of TaKaRa Taq HS (#R007A, TAKARA). Second PCR was performed with a total volume of 50 μl containing 1 μl of PCR product obtained after the first round, 0.4 μM of 5’adaptor-specific primer, 0.4 μM of 2^nd^ round gene-specific primer, 200 μM of each dNTP, 1x PCR buffer, and 1.25 units of TaKaRa Taq HS (#R007A, TAKARA). Following conditions were used for both PCRs: preheating at 95°C for 3 min, followed by 30 cycles of denaturation at 94°C for 30 s, annealing at 56°C for 30 s, extension at 72°C for 30 s, and then final extension at 72°C for 7 min.

For 3’ RACE assay, 15 μg of DNA-depleted RNA was incubated with 25 units of calf intestinal alkaline phosphatase (CIP) (#M0290, NEB) in a 50 μl reaction volume for 1 h at 37°C. After incubation, RNA reacted with CIP was purified and 1.25 μM 3’RACE adaptor (5’-phosphate-UUC ACU GUU CUU AGC GGC CGC AUG CUC-idT −3’) was mixed. The samples were denatured at 95°C for 5 min. The adaptor was ligated overnight using T4 RNA Ligase 1 at 17°C. The adaptor-ligated RNA was purified, annealed with gene-specific primer and reverse-transcribed using AffinityScript Multiple Temperature Reverse Transcriptase (#600107, Agilent) with following conditions; 42°C for 20 min, 55°C for 20 min and 70°C for 15 min in a 20 μl reaction. Reverse-transcribed cDNA of 1 μl was amplified by PCR in a total volume of 50 μl containing 0.4 μM of 3’adaptor-specific primer, 0.4 μM of gene-specific primer, 200 μM of each dNTP, 1x PCR buffer, and 1.25 units of TaKaRa Taq HS (#R007A, TAKARA). The PCR was conducted as follows: preheating at 95°C for 3 min, followed by 30 cycles of denaturation at 94°C for 30 s, annealing at 56°C for 30 s, extension at 72°C for 30 s, and then final extension at 72°C for 7 min.

PCR products were separated on 4% NuSieve GTG agarose gel, the band of interest was cut, gel-eluted, and cloned into a BamHI/HindIII digested pUC19. Bacterial colonies obtained after the transformation were screened by colony PCR. The PCR fragments were purified by QIAquick PCR purification kit and sequenced with an ABI Genetic analyzer 3500 (Applied Biosystems).

### Northern blotting

Purified RNA of 10 μg was denatured at 95°C for 5 min in gel loading buffer II (#AM8547, Invitrogen) and separated by 6% polyacrylamide gel with 7M urea at 300 V for 2 h. Separated RNA was electro-transferred from the gel to Hybond-N+ membranes (GE Healthcare) at 50 V for 1 h at 4°C and the membranes were UV-crosslinked (120 mJ/cm^2^). Northern blotting was performed with the Roche DIG system. DNA probes for 5S rRNA and SPA0079 were amplified with PCR using primers described in Table S2 and PCR DIG Probe Synthesis Kit (#11636090910, Roche). The reaction was performed with a total volume of 50 μl containing 0.2 μM of forward primer, 0.2 μM of reverse primer, 200 μM of dATP, dCTP, and dGTP, 130 μM of dTTP, 70 μM of DIG-11-dUTP, 1x PCR buffer, and 2.625 units of Enzyme mix, Expand High Fidelity (#1732641, Roche). The PCR conditions were set as follows: preheating at 95°C for 2 min, followed by 30 cycles of denaturation at 95°C for 30 s, annealing at 60°C for 30 s, extension at 72°C for 40 s and then final extension at 72°C for 7 min.

UV-crosslinked membranes were prehybridized with 10 ml of DIG EasyHyb for 30 min at 50°C. Thereafter, DIG-labelled DNA probes were hybridized overnight at 50°C in 15 ml of DIG EasyHyb. Membranes were washed every 15 min in 5×Saline Sodium Citrate (SSC)/0.1% SDS, 1×SSC/0.1% SDS and 0.5× SSC/0.1% SDS buffers at 50°C followed by washing with maleic acid buffer (0.1 M maleic acid, 0.15 M NaCl, 0.3% Tween-20 pH 7.5) for 5 min at 37°C. Thereafter, membranes were blocked with blocking solution (#11585762001, Roche) for 45 min at 37°C, and probed with 75 mU/ml Anti-Digoxigenin-AP (#11093274001, Roche) in blocking solution for 45 min at 37°C. Membranes were then washed again in maleic acid wash buffer in two 15-min steps and equilibrated with detection buffer (0.1 M Tris–HCl, 0.1 M NaCl, pH 9.5). Signals were visualized by CDP-star (#12041677001, Roche) with Fusion Fx Imaging System (Vilber-Lourmat).

### Western blotting

One-hundred bacterial cultures at OD_600_ = 2.0 were centrifuged at 10,000 rpm for 2 min at 22°C and resuspended in 100 μl 1xProtein loading buffer, followed by incubation for 5 min at 95°C. Aliquots of 5 μl were separated on 10% TGX gel (Bio-Rad) and subsequently transferred onto a polyvinylidene difluoride (PVDF) membrane using a Transblot Turbo Transfer System (Bio-Rad).

For the CLIP procedure, 5 μl aliquots of heat-denatured immunoprecipitated RNA-protein complex were loaded and separated via SDS-polyacrylamide gel electrophoresis (12% resolving gel) at 30 mA per gel while moving through the stacking gel, which was then increased to 40 mA. After electrophoresis, proteins were electro-transferred onto a PVDF membrane.

Transferred membranes were blocked in 1xTBS-T buffer (20 mM Tris, 150 mM NaCl, 0.1% Tween20, pH 7.6) with 10% skim milk for 45 min. Thereafter, the membrane was probed overnight at 4°C with monoclonal α-FLAG (#31430, ThermoFisher Scientific; 1:10,000) antibody diluted in 1×TBS-T buffer containing 3% bovine serum albumin, washed thrice for 5 min each in 1×TBS-T buffer, probed for 1 h with anti-mouse-HRP-antibody (F1804, Sigma-Aldrich; 1:50,000) diluted in 1×TBS-T buffer containing 3% bovine serum albumin, and washed thrice for 5 min each in 1×TBS-T buffer. Chemiluminescent signals were detected using Immobilon Western Chemiluminescent HRP substrate (#WBKLS0500, Millipore) and measured with Fusion Fx Imaging System (Vilber-Lourmat).

## Data availability

Raw sequence data are available in the DDBJ Sequenced Read Archive under the accession number DRA010307.

## ACKNOWLEDGEMENTS

We thank Akiko Yokota and Tetsushi Suyama for their assistance with the registration of this study in AIST. We also thank Thorsten Bischler for his help with CLIP-seq analysis and Jörg Vogel for his helpful comments on this study.

This study was supported by Grant-in-Aid for JSPS Research Fellow Grant Number 17J10663. The sequencing of CLIP-seq libraries was supported by the JSPS KAKENHI Grant Number 16H06279 (PAGS).

K.C. designed study, performed experiments and data analysis, acquired funding, and wrote manuscript. L.B. performed data analysis and participated in manuscript writing. K.T. performed experiments. N.N and S.T. participated in manuscript writing and supervision.

---

**Figure S1.**
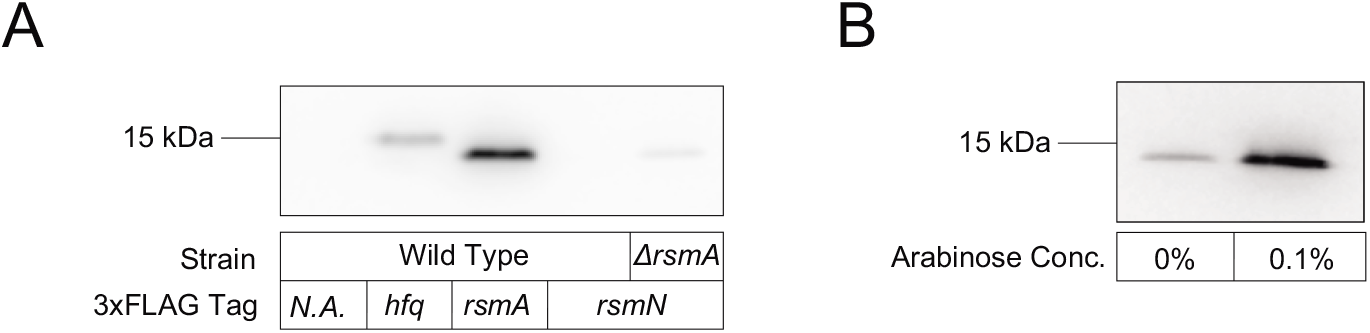
Detection of the 3×FLAG tagged proteins. (A) The presence of the tag was confirmed by evaluating the cell extracts at OD_600_ = 2.0 obtained from PAO1 WT, *hfq*::3×FLAG, *rsmA*::3×FLAG, *rsmN*::3×FLAG, and *rsmN*::3×FLAGΔ*rsmA* strains via western blot analysis using anti-FLAG antibody. (B) The presence of the tag was confirmed by evaluating the cell extracts at OD_600_ = 2.0 obtained from PAO1 WT with pJN-*rsmN*::3×FLAG plasmid via western blot analysis using anti-FLAG antibody. The medium was supplemented with arabinose at 0.1% concentration when OD_600_ reached 0.8.

**Figure S2.**
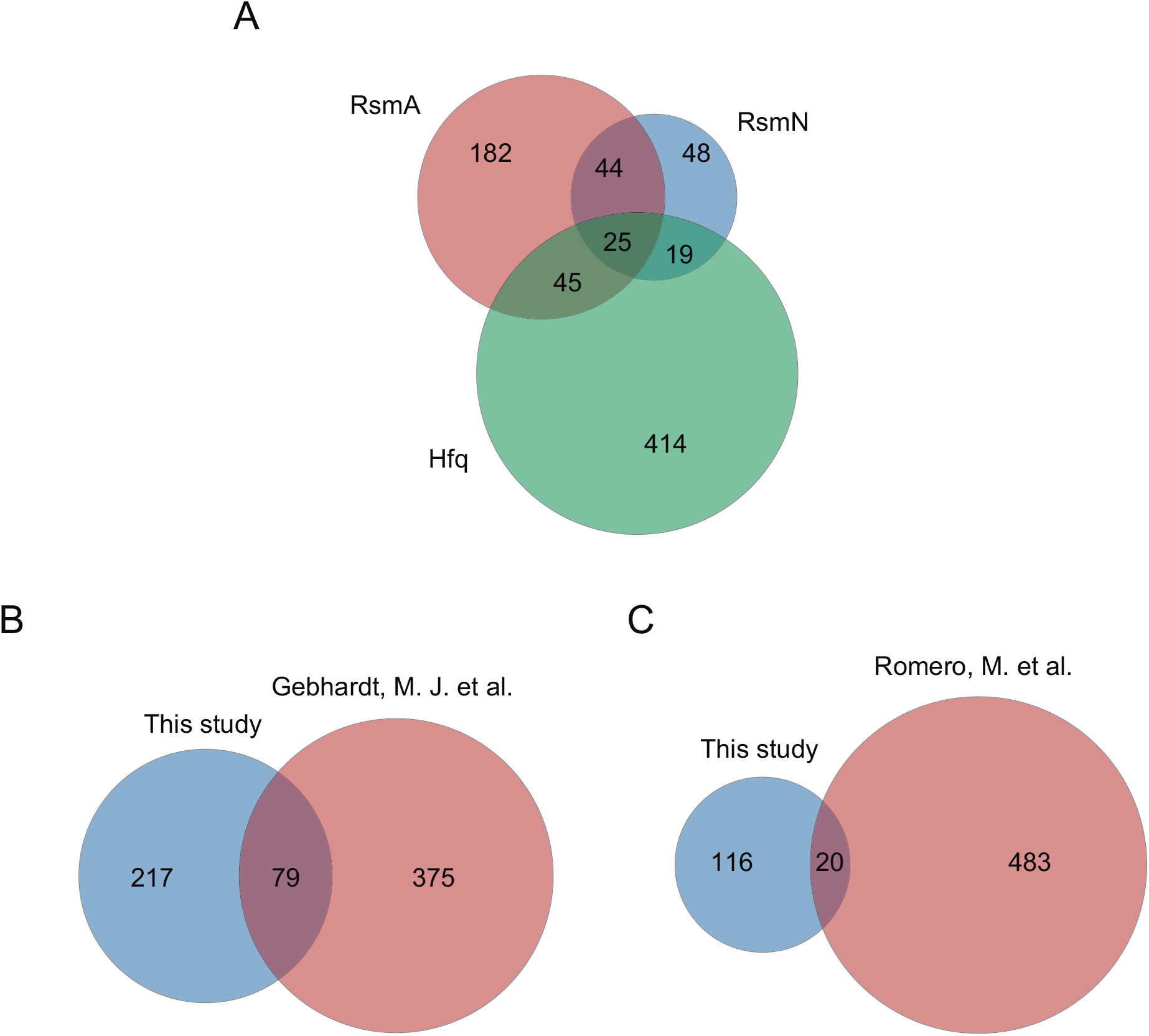
Venn diagram comparing the RsmA/N CLIP-seq to published data. (A) Genes with overlapping peaks between RsmA/N and Hfq. (B) Comparing the genes detected by RsmA CLIP-seq with the list of transcripts detected by ChIPPAR-seq for *P. aeruginosa* RsmA (20). (C) Comparing the genes detected by RsmN CLIP-seq with RsmN-binding genes detected by RIP-seq for *P. aeruginosa* RsmN (19).

**Figure S3.**
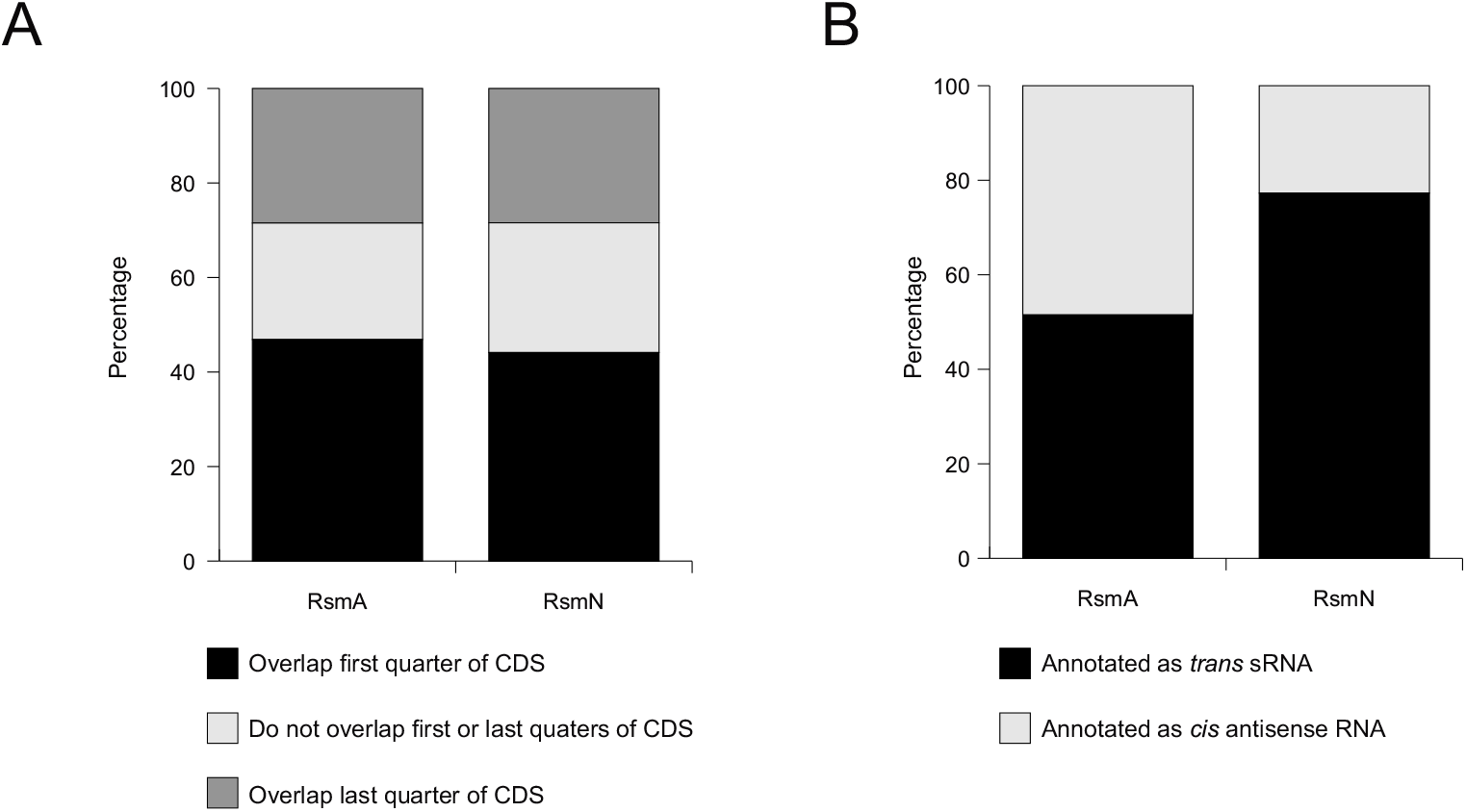
Localization of RsmA/N CLIP-seq peaks relative to known or predicted coding sequences (CDSes) and ncRNAs. (A) Peaks identified in CDSes were classified according to the quadrant positions. (B) Peaks identified in ncRNAs were classified into *trans-sRNAs* and *cis-*antisense RNAs.

**Figure S4.**
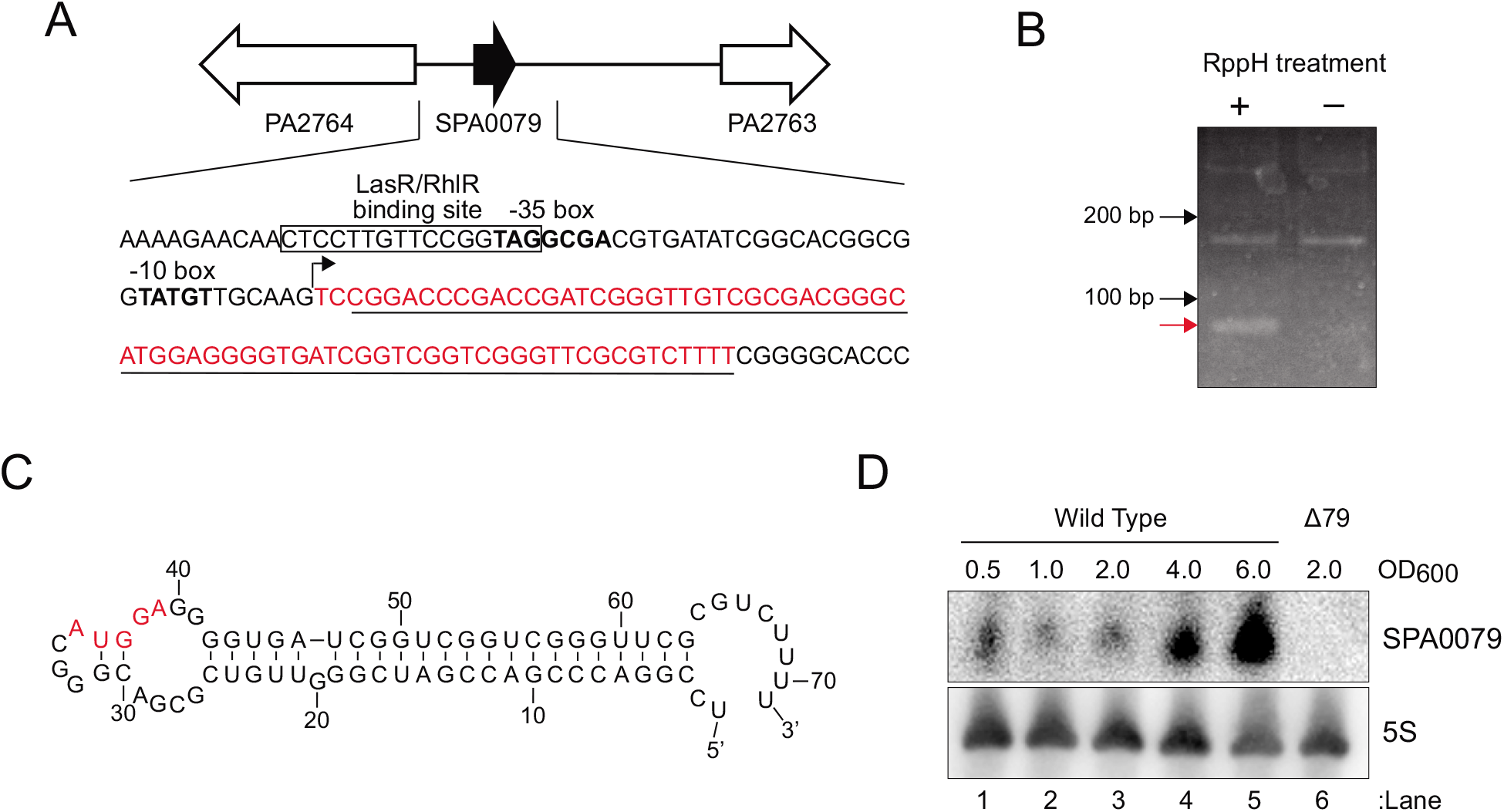
*P. aeruginosa* SPA0079 sRNA. (A) Genetic organization of the SPA0079 sRNA. The −10 and −35 regions of the SPA0079 sRNA promoter are shown in bold. Transcription start site is indicated as an arrow. Underline indicates Hfq-binding site. Red letters indicate the SPA0079 sRNA sequence decided by 5’/3’ RACE. Putative LasR/RhlR binding site is boxed. (B) Representative figure of 5’ RACE of the SPA0079 sRNA. The band corresponding PCR product from 5’ end of the SPA0079 is indicated by red arrow. Non-specific bands detected in both RppH plus and minus conditions were identified as sequences in 23S rRNA. (C) Secondary structure of SPA0079 in *P. aeruginosa* PAO1 predicted by Mfold (34). Red letters indicate AUGGA sequence as a common binding motif of RsmA/N. (D) Northern blot analysis of total RNA isolated from PAO1 wild-type and ΔSPA0079 strain. RNA was extracted at the indicated OD_600_. SPA0079 and 5S rRNA was detected using DIG-labelled probes.

**Figure S5.**
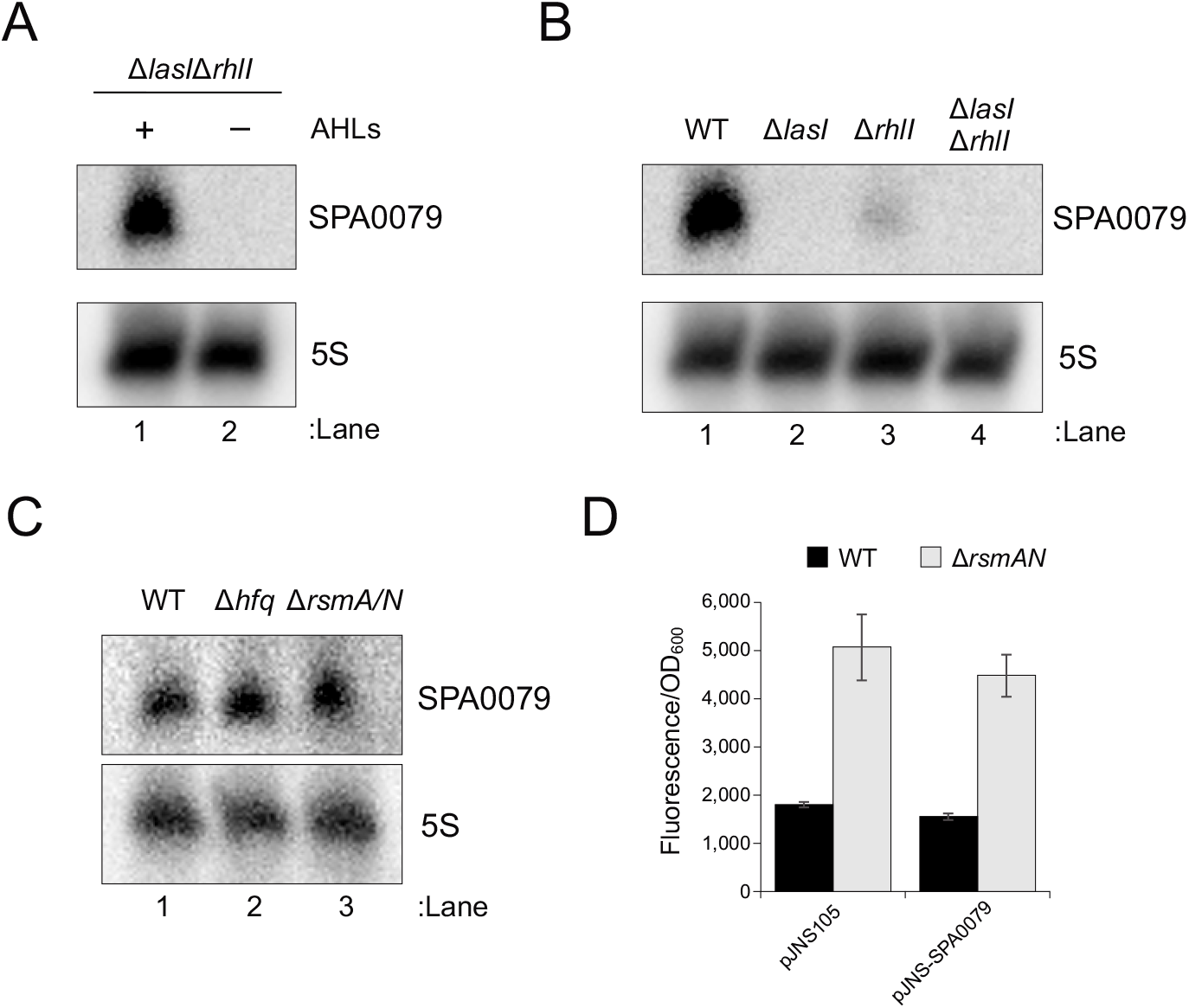
Characterization of the SPA0079 sRNA. (A to C) Northern blot analysis of total RNA isolated from the indicated strains. RNA was extracted at OD_600_ = 6.0. SPA0079 and 5S rRNA was detected using DIG-labelled probes. (D) Super-folder GFP translational fusion assay for *cafA* gene between pJNS105 control vector and pJNS-SPA0079 overexpression vector. For exogenous SPA0079 expression, 0.1% arabinose was added.

**Figure S6.**
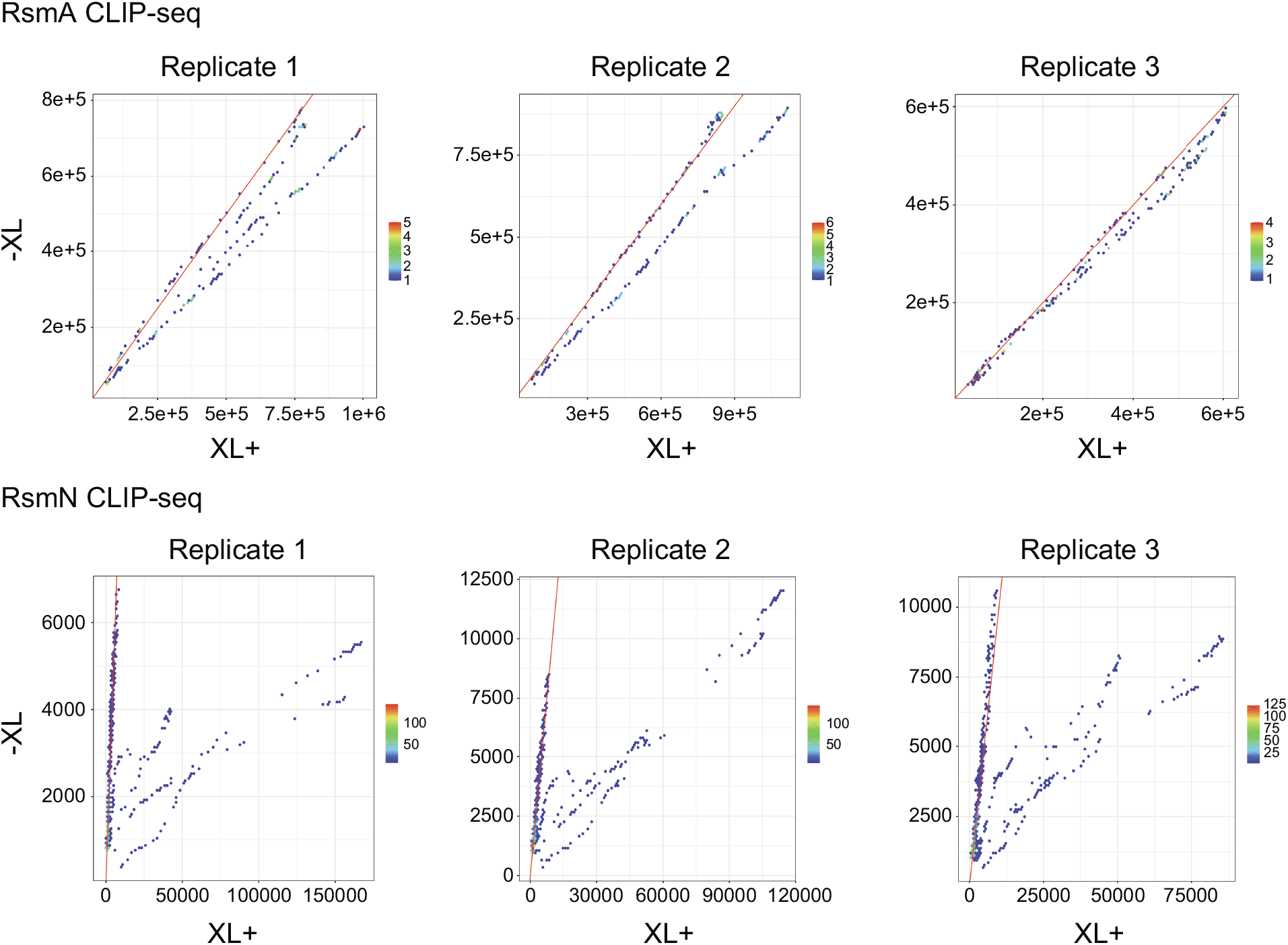
Frequency plots of crosslinked and background samples between each replicate. The axes show read counts and the coloring shows the frequency of each x-y pair. Red line indicates y = x.

Table S1. RsmA/N peaks detected by CLIP-seq

Table S2. The list of strains, plasmids, and oligonucleotides

## REFERENCES

1. Mulcahy LR, Isabella VM, Lewis K. 2014. *Pseudomonas aeruginosa* Biofilms in Disease. Microbial Ecology 68:1–12.

2. Holmqvist E, Wagner EGH. 2017. Impact of bacterial sRNAs in stress responses. Biochem Soc Trans 45:1203–1212.

3. Chao Y, Vogel J. 2010. The role of Hfq in bacterial pathogens. Curr Opin Microbiol 13:24–33.

4. Sonnleitner E, Hagens S, Rosenau F, Wilhelm S, Habel A, Jäger KE, Bläsi U. 2003. Reduced virulence of a *hfq* mutant of *Pseudomonas aeruginosa* O1. Microbial Pathogenesis 35:217–228.

5. Sonnleitner E, Schuster M, Sorger-Domenigg T, Greenberg EP, Blasi U. 2006. Hfq-dependent alterations of the transcriptome profile and effects on quorum sensing in *Pseudomonas aeruginosa*. Mol Microbiol 59:1542–1558.

6. Pusic P, Sonnleitner E, Krennmayr B, Heitzinger DA, Wolfinger MT, Resch A, Bläsi U. 2018. Harnessing metabolic regulation to increase Hfq-dependent antibiotic susceptibility in *Pseudomonas aeruginosa*. Front Microbiol 9:2709.

7. Romeo T, Gong M, Liu MY, Brun-Zinkernagel AM. 1993. Identification and molecular characterization of *csrA*, a pleiotropic gene from *Escherichia coli* that affects glycogen biosynthesis, gluconeogenesis, cell size, and surface properties. J Bacteriol 175:4744–4755.

8. Babitzke P, Romeo T. 2007. CsrB sRNA family: sequestration of RNA-binding regulatory proteins. Curr Opin Microbiol 10:156–163.

9. Holmqvist E, Vogel J. 2013. A small RNA serving both the Hfq and CsrA regulons. Genes Dev 27:1073–1078.

10. Kay E, Humair B, Denervaud V, Riedel K, Spahr S, Eberl L, Valverde C, Haas D. 2006. Two GacA-dependent small RNAs modulate the quorum-sensing response in *Pseudomonas aeruginosa*. J Bacteriol 188:6026–6033.

11. Miller CL, Romero M, Karna SL, Chen T, Heeb S, Leung KP. 2016. RsmW, *Pseudomonas aeruginosa* small non-coding RsmA-binding RNA upregulated in biofilm versus planktonic growth conditions. BMC Microbiol 16:155.

12. Janssen KH, Diaz MR, Gode CJ, Wolfgang MC, Yahr TL. 2018. RsmV, a small noncoding regulatory RNA in *Pseudomonas aeruginosa* that sequesters RsmA and RsmF from target mRNAs. J Bacteriol 200:e00277–18.

13. Morris ER, Hall G, Li C, Heeb S, Kulkarni RV, Lovelock L, Silistre H, Messina M, Camara M, Emsley J, Williams P, Searle MS. 2013. Structural rearrangement in an RsmA/CsrA ortholog of *Pseudomonas aeruginosa* creates a dimeric RNA-binding protein, RsmN. Structure 21:1659–1671.

14. Goodman AL, Merighi M, Hyodo M, Ventre I, Filloux A, Lory S. 2009. Direct interaction between sensor kinase proteins mediates acute and chronic disease phenotypes in a bacterial pathogen. Genes Dev 23:249–259.

15. Brencic A, McFarland KA, McManus HR, Castang S, Mogno I, Dove SL, Lory S. 2009. The GacS/GacA signal transduction system of *Pseudomonas aeruginosa* acts exclusively through its control over the transcription of the RsmY and RsmZ regulatory small RNAs. Mol Microbiol 73:434–445.

16. Irie Y, Starkey M, Edwards AN, Wozniak DJ, Romeo T, Parsek MR. 2010. *Pseudomonas aeruginosa* biofilm matrix polysaccharide Psl is regulated transcriptionally by RpoS and post-transcriptionally by RsmA. Mol Microbiol 78:158–172.

17. Allsopp LP, Wood TE, Howard SA, Maggiorelli F, Nolan LM, Wettstadt S, Filloux A. 2017. RsmA and AmrZ orchestrate the assembly of all three type VI secretion systems in *Pseudomonas aeruginosa*. Proc Natl Acad Sci U S A 114:7707–7712.

18. Heurlier K, Williams F, Heeb S, Dormond C, Pessi G, Singer D, Camara M, Williams P, Haas D. 2004. Positive control of swarming, rhamnolipid synthesis, and lipase production by the posttranscriptional RsmA/RsmZ system in *Pseudomonas aeruginosa* PAO1. J Bacteriol 186:2936–2945.

19. Romero M, Silistre H, Lovelock L, Wright VJ, Chan KG, Hong KW, Williams P, Camara M, Heeb S. 2018. Genome-wide mapping of the RNA targets of the *Pseudomonas aeruginosa* riboregulatory protein RsmN. Nucleic Acids Res 46:6823–6840.

20. Gebhardt MJ, Kambara TK, Ramsey KM, Dove SL. 2020. Widespread targeting of nascent transcripts by RsmA in *Pseudomonas aeruginosa*. Proc Natl Acad Sci U S A 117:10520–10529.

21. Marden JN, Diaz MR, Walton WG, Gode CJ, Betts L, Urbanowski ML, Redinbo MR, Yahr TL, Wolfgang MC. 2013. An unusual CsrA family member operates in series with RsmA to amplify posttranscriptional responses in *Pseudomonas aeruginosa*. Proc Natl Acad Sci U S A 110:15055–15060.

22. Chihara K, Bischler T, Barquist L, Monzon AV, Noda N, Vogel J, Tsuneda S. 2019. Conditional Hfq association with small noncoding RNAs in *Pseudomonas aeruginosa* revealed through comparative UV cross-linking immunoprecipitation followed by high-throughput sequencing. mSystems 4:e00590–19.

23. Kingsford CL, Ayanbule K, Salzberg SL. 2007. Rapid, accurate, computational discovery of Rho-independent transcription terminators illuminates their relationship to DNA uptake. Genome Biol 8:R22.

24. Gill EE, Chan LS, Winsor GL, Dobson N, Lo R, Ho Sui SJ, Dhillon BK, Taylor PK, Shrestha R, Spencer C, Hancock REW, Unrau PJ, Brinkman FSL. 2018. High-throughput detection of RNA processing in bacteria. BMC Genomics 19:223.

25. Ferrara S, Brugnoli M, De Bonis A, Righetti F, Delvillani F, Deho G, Horner D, Briani F, Bertoni G. 2012. Comparative profiling of *Pseudomonas aeruginosa* strains reveals differential expression of novel unique and conserved small RNAs. PLoS One 7:e36553.

26. Gomez-Lozano M, Marvig RL, Molin S, Long KS. 2012. Genome-wide identification of novel small RNAs in *Pseudomonas aeruginosa*. Environ Microbiol 14:2006–2016.

27. Gomez-Lozano M, Marvig RL, Tulstrup MV, Molin S. 2014. Expression of antisense small RNAs in response to stress in *Pseudomonas aeruginosa*. BMC Genomics 15:783.

28. Schulmeyer KH, Diaz MR, Bair TB, Sanders W, Gode CJ, Laederach A, Wolfgang MC, Yahr TL. 2016. Primary and secondary sequence structure requirements for recognition and discrimination of target RNAs by *Pseudomonas aeruginosa* RsmA and RsmF. J Bacteriol 198:2458–2469.

29. Sorger-Domenigg T, Sonnleitner E, Kaberdin VR, Blasi U. 2007. Distinct and overlapping binding sites of *Pseudomonas aeruginosa* Hfq and RsmA proteins on the non-coding RNA RsmY. Biochem Biophys Res Commun 352:769–773.

30. Rocchetta HL, Burrows LL, Lam JS. 1999. Genetics of O-antigen biosynthesis in *Pseudomonas aeruginosa*. Microbiol Mol Biol Rev 63:523–553.

31. Bordi C, Lamy MC, Ventre I, Termine E, Hachani A, Fillet S, Roche B, Bleves S, Mejean V, Lazdunski A, Filloux A. 2010. Regulatory RNAs and the HptB/RetS signalling pathways fine-tune *Pseudomonas aeruginosa* pathogenesis. Mol Microbiol 76:1427–1443.

32. Petrova OE, Sauer K. 2010. The novel two-component regulatory system BfiSR regulates biofilm development by controlling the small RNA *rsmZ* through CafA. J Bacteriol 192:5275–5288.

33. Bouillet S, Ba M, Houot L, Iobbi-Nivol C, Bordi C. 2019. Connected partner-switches control the life style of *Pseudomonas aeruginosa* through RpoS regulation. Sci Rep 9:6496.

34. Zuker M. 2003. Mfold web server for nucleic acid folding and hybridization prediction. Nucleic Acids Res 31:3406–3415.

35. Thomason MK, Voichek M, Dar D, Addis V, FitzGerald DJ, Gottesman S, Sorek R, Greenberg EP, Buchrieser C. 2019. A *rhlI* 5’ UTR-derived sRNA regulates RhlR-dependent quorum sensing in *Pseudomonas aeruginosa*. mBio 10:43.

36. Tree JJ, Granneman S, McAteer SP, Tollervey D, Gally DL. 2014. Identification of bacteriophage-encoded anti-sRNAs in pathogenic *Escherichia coli*. Mol Cell 55:199–213.

37. Holmqvist E, Wright PR, Li L, Bischler T, Barquist L, Reinhardt R, Backofen R, Vogel J. 2016. Global RNA recognition patterns of post-transcriptional regulators Hfq and CsrA revealed by UV crosslinking *in vivo*. EMBO J 35:991–1011.

38. Potts AH, Vakulskas CA, Pannuri A, Yakhnin H, Babitzke P, Romeo T. 2017. Global role of the bacterial post-transcriptional regulator CsrA revealed by integrated transcriptomics. Nat Commun 8:1596.

39. Lovelock L. 2012. RsmN: a new atypical RsmA homologue in Pseudomonas aeruginosa. PhD thesis. Univ of Nottingham, Nottingham, United Kingdom.

40. Lee FCY, Ule J. 2018. Advances in CLIP technologies for studies of protein-RNA interactions. Mol Cell 69:354–369.

41. Van Nostrand EL, Pratt GA, Shishkin AA, Gelboin-Burkhart C, Fang MY, Sundararaman B, Blue SM, Nguyen TB, Surka C, Elkins K, Stanton R, Rigo F, Guttman M, Yeo GW. 2016. Robust transcriptome-wide discovery of RNA-binding protein binding sites with enhanced CLIP (eCLIP). Nat Methods 13:508–514.

42. Maticzka D, Ilik IA, Aktas T, Backofen R, Akhtar A. 2018. uvCLAP is a fast and non-radioactive method to identify *in vivo* targets of RNA-binding proteins. Nat Commun 9:1142.

43. Ilik IA, Aktas T, Maticzka D, Backofen R, Akhtar A. 2019. FLASH: ultra-fast protocol to identify RNA-protein interactions in cells. Nucleic Acids Res 48:e15.

44. Müller P, Gimpel M, Wildenhain T, Brantl S. 2019. A new role for CsrA: promotion of complex formation between an sRNA and its mRNA target in *Bacillus subtilis*. RNA Biol:1–16.

45. Olejniczak M, Storz G. 2017. ProQ/FinO-domain proteins: another ubiquitous family of RNA matchmakers? Mol Microbiol 104:905–915.

46. van Biesen T, Frost LS. 1994. The FinO protein of IncF plasmids binds FinP antisense RNA and its target, *traJ* mRNA, and promotes duplex formation. Mol Microbiol 14:427–436.

47. Smirnov A, Forstner KU, Holmqvist E, Otto A, Gunster R, Becher D, Reinhardt R, Vogel J. 2016. Grad-seq guides the discovery of ProQ as a major small RNA-binding protein. Proc Natl Acad Sci U S A 113:11591–11596.

48. Smirnov A, Wang C, Drewry LL, Vogel J. 2017. Molecular mechanism of mRNA repression *in trans* by a ProQ-dependent small RNA. EMBO J 36:1029–1045.

49. Melamed S, Adams PP, Zhang A, Zhang H, Storz G. 2019. RNA-RNA interactomes of ProQ and Hfq reveal overlapping and competing roles. Mol Cell 77:411–425.e7.

50. Ross JA, Ellis MJ, Hossain S, Haniford DB. 2013. Hfq restructures RNA-IN and RNA-OUT and facilitates antisense pairing in the Tn10/IS10 system. RNA 19:670–684.

51. Pessi G, Williams F, Hindle Z, Heurlier K, Holden MT, Camara M, Haas D, Williams P. 2001. The global posttranscriptional regulator RsmA modulates production of virulence determinants and N-acylhomoserine lactones in *Pseudomonas aeruginosa*. J Bacteriol 183:6676–6683.

52. Duss O, Michel E, Diarra dit Konte N, Schubert M, Allain FH. 2014. Molecular basis for the wide range of affinity found in Csr/Rsm protein-RNA recognition. Nucleic Acids Res 42:5332–5346.

53. Burrows LL, Chow D, Lam JS. 1997. *Pseudomonas aeruginosa* B-band O-antigen chain length is modulated by Wzz (Ro1). J Bacteriol 179:1482–1489.

54. Deretic V, Schurr MJ, Yu H. 1995. *Pseudomonas aeruginosa*, mucoidy and the chronic infection phenotype in cystic fibrosis. Trends in Microbiology 3:351–356.

55. Cross AR, Goldberg JB. 2019. Remodeling of O antigen in mucoid *Pseudomonas aeruginosa* via transcriptional repression of *wzz2*. mBio 10:e02914–18.

56. Stacey SD, Williams DA, Pritchett CL. 2017. The *Pseudomonas aeruginosa* two-component regulator AlgR directly activates *rsmA* expression in a phosphorylation independent manner. J Bacteriol 199:e00048–17.

57. Ghafoor A, Hay ID, Rehm BH. 2011. Role of exopolysaccharides in *Pseudomonas aeruginosa* biofilm formation and architecture. Appl Environ Microbiol 77:5238–5246.

58. Augustin DK, Song Y, Baek MS, Sawa Y, Singh G, Taylor B, Rubio-Mills A, Flanagan JL, Wiener-Kronish JP, Lynch SV. 2007. Presence or absence of lipopolysaccharide O antigens affects type III secretion by *Pseudomonas aeruginosa*. J Bacteriol 189:2203–2209.

59. Huang H, Shao X, Xie Y, Wang T, Zhang Y, Wang X, Deng X. 2019. An integrated genomic regulatory network of virulence-related transcriptional factors in *Pseudomonas aeruginosa*. Nat Commun 10:2931.

60. Intile PJ, Diaz MR, Urbanowski ML, Wolfgang MC, Yahr TL. 2014. The AlgZR two-component system recalibrates the RsmAYZ posttranscriptional regulatory system to inhibit expression of the *Pseudomonas aeruginosa* type III secretion system. J Bacteriol 196:357–366.

61. Wang Z, Huang X, Liu Y, Yang G, Liu Y, Zhang X. 2017. GacS/GacA activates pyoluteorin biosynthesis through Gac/Rsm-RsmE cascade and RsmA/RsmE-driven feedback loop in *Pseudomonas protegens* H78. Mol Microbiol 105:968–985.

62. Elowitz MB, Leibler S. 2000. A synthetic oscillatory network of transcriptional regulators. Nature 403:335–338.

63. Chen R, Weng Y, Zhu F, Jin Y, Liu C, Pan X, Xia B, Cheng Z, Jin S, Wu W. 2016. Polynucleotide phosphorylase regulates multiple virulence factors and the stabilities of small RNAs RsmY/Z in *Pseudomonas aeruginosa*. Front Microbiol 7:247.

64. Janssen KH, Corley JM, Djapgne L, Cribbs JT, Voelker D, Slusher Z, Nordell R, Regulski EE, Kazmierczak BI, McMackin EW, Yahr TL. 2020. Hfq and sRNA 179 inhibit expression of the *Pseudomonas aeruginosa* cAMP-Vfr and type III secretion regulons. mBio 11:e00363–20.

65. Hmelo LR, Borlee BR, Almblad H, Love ME, Randall TE, Tseng BS, Lin C, Irie Y, Storek KM, Yang JJ, Siehnel RJ, Howell PL, Singh PK, Tolker-Nielsen T, Parsek MR, Schweizer HP, Harrison JJ. 2015. Precision-engineering the *Pseudomonas aeruginosa* genome with two-step allelic exchange. Nat Protoc 10:1820–1841.

66. Maseda H, Sawada I, Saito K, Uchiyama H, Nakae T, Nomura N. 2004. Enhancement of the *mexAB-oprM* efflux pump expression by a quorum-sensing autoinducer and its cancellation by a regulator, MexT, of the *mexEF-oprN* efflux pump operon in *Pseudomonas aeruginosa*. Antimicrob Agents Chemother 48:1320–1328.

67. Martin M. 2011. Cutadapt removes adapter sequences from high-throughput sequencing reads. EMBnetjournal 17.

68. Xu H, Luo X, Qian J, Pang X, Song J, Qian G, Chen J, Chen S. 2012. FastUniq: a fast *de novo* duplicates removal tool for paired short reads. PLoS One 7:e52249.

69. Forstner KU, Vogel J, Sharma CM. 2014. READemption-a tool for the computational analysis of deep-sequencing-based transcriptome data. Bioinformatics 30:3421–3423.

70. Hoffmann S, Otto C, Doose G, Tanzer A, Langenberger D, Christ S, Kunz M, Holdt LM, Teupser D, Hackermuller J, Stadler PF. 2014. A multi-split mapping algorithm for circular RNA, splicing, trans-splicing and fusion detection. Genome Biol 15:R34.

71. Holmqvist E, Li L, Bischler T, Barquist L, Vogel J. 2018. Global maps of ProQ binding *in vivo* reveal target recognition via RNA structure and stability control at mRNA 3’ ends. Mol Cell 70:971–982 e6.

72. Anders S, Huber W. 2010. Differential expression analysis for sequence count data. Genome Biol 11:R106.

73. Langenberger D, Bermudez-Santana C, Hertel J, Hoffmann S, Khaitovich P, Stadler PF. 2009. Evidence for human microRNA-offset RNAs in small RNA sequencing data. Bioinformatics 25:2298–2301.

74. Love MI, Huber W, Anders S. 2014. Moderated estimation of fold change and dispersion for RNA-seq data with DESeq2. Genome Biol 15:550.

75. Bailey TL, Johnson J, Grant CE, Noble WS. 2015. The MEME Suite. Nucleic Acids Res 43:W39–49.

76. Yao Z, Weinberg Z, Ruzzo WL. 2006. CMfinder--a covariance model based RNA motif finding algorithm. Bioinformatics 22:445–452.

77. Weinberg Z, Breaker RR. 2011. R2R--software to speed the depiction of aesthetic consensus RNA secondary structures. BMC Bioinformatics 12:3.

78. Corcoran CP, Podkaminski D, Papenfort K, Urban JH, Hinton JC, Vogel J. 2012. Superfolder GFP reporters validate diverse new mRNA targets of the classic porin regulator, MicF RNA. Mol Microbiol 84:428–445.

79. Wilton R, Ahrendt AJ, Shinde S, Sholto-Douglas DJ, Johnson JL, Brennan MB, Kemner KM. 2017. A new suite of plasmid vectors for fluorescence-based imaging of root colonizing *Pseudomonads*. Front Plant Sci 8:2242.

80. Beaudoin T, Zhang L, Hinz AJ, Parr CJ, Mah TF. 2012. The biofilm-specific antibiotic resistance gene *ndvB* is important for expression of ethanol oxidation genes in *Pseudomonas aeruginosa* biofilms. J Bacteriol 194:3128–3136.

81. Livak KJ, Schmittgen TD. 2001. Analysis of relative gene expression data using real-time quantitative PCR and the 2(-Delta Delta C(T)) Method. Methods 25:402–408.

82. Argaman L, Hershberg R, Vogel J, Bejerano G, Wagner EGH, Margalit H, Altuvia S. 2001. Novel small RNA-encoding genes in the intergenic regions of *Escherichia coli*. Curr Biol 11:941–950.

